# Dynamic Functional Pathway Development in Type 1 Spinal Interneurons: Stage-specific roles of retinoic acid activity

**DOI:** 10.1101/2025.07.29.667367

**Authors:** Dina Rekler, Sarah Kagan, Noa Rachel Krutous, Susanna Ventriglia, Gilgi Friedlander, Chaya Kalcheim

## Abstract

**Background:** dI1 spinal interneuron development requires precise coordination of proliferation, differentiation, and migration, yet the molecular and cellular properties underlying the temporal coupling of these processes remain unresolved.

**Results:** Single-cell RNA sequencing of the E4 quail neural tube under control and retinoic acid (RA)-deprived conditions generated a high-resolution atlas spanning a continuous trajectory from dorsal progenitors to differentiated neurons. Pseudotime and RNA velocity analyses revealed transitions consistent with medial-to-lateral displacement in vivo. Early progenitors expressed cell cycle and BMP-associated programs, followed by induction of cytoskeletal and extracellular matrix remodelling genes and later activation of pan-neuronal and synaptic pathways. BMP signaling declined during differentiation through upregulation of BMP antagonists, whereas apoptosis-related genes were poised during specification without overt cell death. Autophagy-associated genes peaked during differentiation. Disruption of RA signaling by dominant-negative RARα403 uncovered stage-specific functions of RA. In progenitors, RA inhibition increased proliferation. In differentiating neurons, it upregulated BARHL1/2 and enhanced ion channel, synaptic, cytoskeletal, and extracellular matrix gene programs. RA inhibition also impaired ventral displacement of BarHL2-positive dI1 neurons, causing abnormal positioning.

**Conclusions:** These findings define a dI1-centered single-cell atlas of the progenitor-to-neuron transition and identify stage-specific roles for RA in dorsal hinterneuron maturation.

**Key findings:**

- We provide a high-resolution single-cell atlas of dI1 spinal interneuron development in the E4 quail neural tube, analysis of this single stage reveals a continuous trajectory from proliferative dorsal progenitors to differentiated neurons.
- Development along this trajectory is tightly staged: early cells are dominated by cell-cycle and progenitor-maintenance programs, while later cells progressively activate cytoskeletal, extracellular matrix, axon-guidance, synaptic, and mature neuronal pathways.
- BMP signaling is strongest in early dI1 progenitors and is progressively attenuated during differentiation through the induction of BMP inhibitors, supporting a stage-specific role for BMP in early specification rather than late maturation.
- Retinoic acid acts in a stage-dependent manner: RA inhibition increases progenitor proliferation, alters differentiation-associated gene programs, and upregulates neuronal maturation, ion-channel, and synaptic genes in differentiating neurons.
- Repressing RA signaling causes BarHL2-positive dI1 neurons to fail their normal ventral migration. Instead, they accumulate near their dorsal origin and show ectopic localization, indicating that RA is required for proper migration.

## Introduction

Neural development transforms a seemingly uniform neuroepithelium into a diverse array of neuronal and glial cell types that arise in appropriate numbers, positions and times. How closely related progenitors generate such diversity through spatially and temporally patterned signaling and transcriptional networks remains a central question in developmental biology ^1–4^

The developing spinal cord is a particularly tractable model for addressing this problem, as discrete classes of interneurons and motoneurons are organized along the dorsoventral axis and contribute to well-defined sensorimotor circuits. Its dorsal region primarily consists of sensory interneurons that process and relay sensory information, classified into six discrete subpopulations, from dI1-to dI6 ^5–11^

We focus on dI1 interneurons, which relay proprioceptive signals from the periphery to higher brain centers^12–14^. These glutamatergic neurons initially project axons commissurally across the floor plate at E4 in chick, but from E4.5 onward, there is a progressive shift to ipsilateral projections, with most becoming ipsilateral by late embryonic stages, a pattern conserved in mouse ^15–17^. They are among the earliest-born dorsal interneurons, arising from the dorsal-most progenitor domain (dp1) ^18,19^. Their specification depends on the roof plate (RP), as its genetic ablation eliminates dI1 interneurons ^20^, and is tightly regulated by RP-specific factors such as BMPs ^18,21–23^, initially secreted by the dorsal neural tube (NT) and later by the definitive RP ^24–26^. BMPs were identified as dorsalizing factors that induce expression of the proneural gene *ATOH1*, which marks the dp1 domain and initiates dI1 neurogenesis ^19,27–29^. BMP4 further upregulates Wnt1 and 3 ^30^ and acts upstream of Wnt/β-catenin signals to control expression of determinants such as *OLIG3* which marks the dI1-3 subsets ^31^. Work from several groups suggested that Wnt signals contribute to controling proliferation within dorsal progenitor domains, including dp1, in addition to their roles in fate specification ^6,32,33^. Next, dI1 interneurons migrate ventrally to distinct locations and integrate into specific circuits. The molecular cues guiding this migration and subsequent diversification are incompletely characterized ^8,34^

Retinoic acid (RA) adds an additional layer of regulation in the dorsal spinal cord. It is first produced in the paraxial mesoderm, and later in the developing RP close to the conclusion of neural crest (NC) formation. At this stage, RA contributes to the termination of NC production by repressing BMP and Wnt activities ^26,35^, both of which are required for the onset of NC emigration ^24,30,36^. In absence of RA signaling, NC progenitors keep exiting the NT well into the RP stage, and the temporal segregation between NC and ensuing RP traits is lost ^9,26^. At this stage, the RP region contains NC, RP and dI1 cells, and single-cell RNA-seq analysis revealed progenitors co-expressing markers of all three lineages ^9^. Moreover, absence of RA or gain of Notch activity, disrupts the boundary that normally separates RP from dI1 cells and rescue experiments suggest that RA is required for RP/dI1 segregation by restricting Notch signaling ^9,37^.

Thus, RP-derived RA is necessary for the separation of lineages in both time (NC/RP) and space (RP/dI1). Whereas the roles of RA in lineage segregation have begun to be clarified, its possible function/s on various stages of dI1 development remain unexplored. Resolving this could unveil new layers of spatiotemporal control leading to the assembly of neural circuits.

Here, we used scRNA-seq to map dI1 development in the quail embryo at E4, a critical phase characterized by the simultaneous presence of, and transition from proliferative progenitors to differentiated neurons. The quail embryo provides an accessible model in which E4 corresponds to early neurogenic stages when dorsal progenitors are transitioning toward post-mitotic interneurons. Within this single-timepoint dataset, we apply trajectory inference to organize dI1 cells along a continuous pseudotime axis and define stage-associated transcriptional modules. Whereas early phases are enriched for proliferation and progenitor maintenance programs, later phases show sequential induction of cytoskeletal, extracellular matrix and neuronal effector genes. BMP signaling is prominent in early progenitors and appears progressively attenuated as differentiation proceeds. Apoptosis-related genes are primed without overt cell death, autophagy genes increase during differentiation, and neuronal identity and function genes are activated late as cells acquire mature properties.

We further compare control embryos with embryos subjected to RA inhibition. The latter enhances cell proliferation and differentiation, and modulates ventral localization of dI1 neurons. Together, our study provides a dI1-centered atlas in the quail spinal cord and highlights candidate RA-linked programs that may coordinate progenitor patterning and neuronal maturation during dorsal interneuron development.

## Results

### Uncovering the dynamics of normal dI1 interneuron development

#### A single-cell atlas of dI1 interneuron development in the quail NT

To resolve dI1 developmental states within a defined temporal window, we analyzed our previously generated single cell RNA-seq dataset from dorsal E4 quail NT (Fig. 1A) ^9^. At this stage, the NT contained both proliferative progenitors of the various dorsal lineages primarily located at the main body of the UMAP, and also differentiating post-mitotic neurons distributed along the distinct arms (Fig. 1B). Within this framework, dI1 interneurons were prominently represented in four clusters (Fig. 1A,C), defined by characteristic genes and gene combinations (Fig. 1D; Sup. Figs. 1–4), providing a snapshot of their developmental progression at this critical stage.

**Figure 1.**
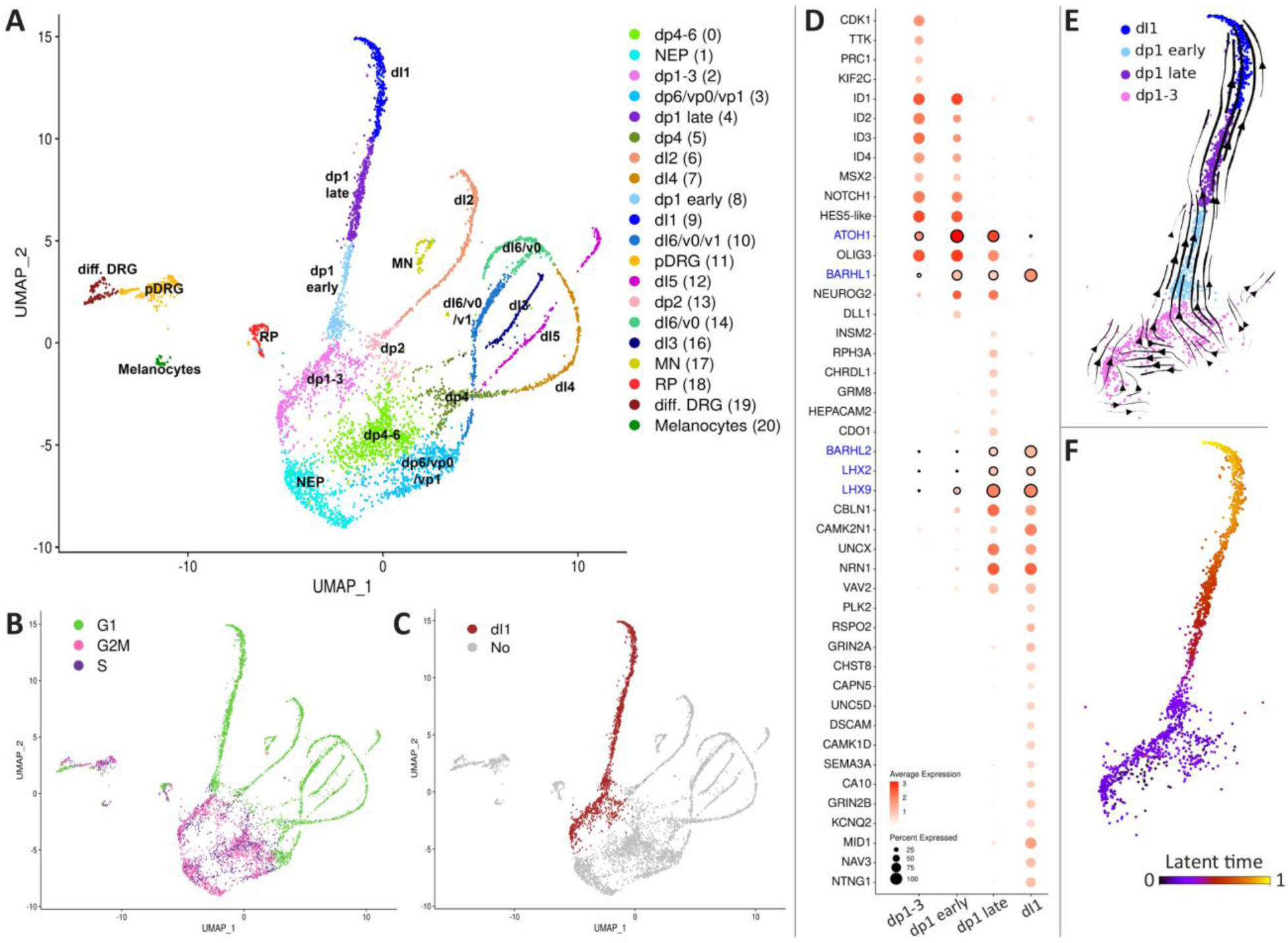
scRNA-seq delineates progressive phases of dI1 interneuron maturation in the E4 quail embryo. A) UMAP of the control sample, colored by cluster identity. Cluster numbers indicated in parentheses. B) UMAP colored by cell cycle phase. C) UMAP of the control sample. Cells belonging to clusters associated with dI1 interneurons are shown in red. D) Dot plot visualizing selected marker gene expression in each cluster belonging to the dI1 trajectory. The size of the dot corresponds to the percentage of cells expressing the gene in each cluster. The color represents the average expression level. In blue, canonical genes of the dI1 lineage. Genes were initially identified by differential expression analysis (linear fold change > 1.4, adjusted p-value < 0.05). E) RNA velocity vectors projected on a UMAP of the dI1 clusters. F) Latent time analysis of the dI1 clusters. The earliest point is depicted in black/purple and the latest in yellow.

The dp1–3 cluster uniquely expressed *CDK1, TTK, PRC1* and *KIF2C*, consistent with a shared proliferative state of the three dorsal-most progenitor types (Fig. 1D; Sup. Fig. 1). The dp1-early cluster showed increased expression of dI1 lineage markers *ATOH1* and *BARHL1*, broader dorsal determinants *OLIG3* and *NEUROG2*, and uniquely expressed *DLL1*, indicative of emerging neuronal patterning, while retaining progenitor-associated markers shared with early dp1–3 cells (*ID1–4, MSX2, NOTCH1, HES5-LIKE*). Late dp1 progenitors upregulated neuronal differentiation factors *LHX2*/9 and *BARHL2* and further increased *BARHL1*, together with neuronally enriched genes such as *CBLN1, CAMK2N1* and *NRN1*, yet still expressed *ATOH1, OLIG3* and *NEUROG2*, consistent with an intermediate dI1 neuroblast state. The dI1 cluster displayed high *BARHL1/2* and *LHX2/9* ^5,6,38^, as well as *RSPO2, GRIN2A, SEMA3A, CAPN5, GRIA1* and others, compatible with neuronal differentiation and onset of axon guidance programs, and shared many pan-neuronal genes with other dorsal interneurons (e.g. *ACTL6B, CAMK2N1, CELF4, EBF1, NSG2, NRXN1, GAP43*), indicating convergence toward a mature neuronal profile. Together, the distinct and partially overlapping expression profiles across these four clusters define a transcriptional continuum from cycling dp1–3 progenitors through intermediate precursors to differentiated dI1 neurons in the E4 dorsal NT. Importantly, in all clusters, numerous markers with so far uncharacterized roles in dI1 development were identified (e.g; *PLCL1, PKIG, LIFR, INSM2, TMEM163, MAB2IL2, GRIN2A, PLK, CHST8,* etc) (Sup. Figs. 1-4).

Trajectory inference by RNA velocity (Fig. 1E) and latent time analysis (Fig. 1F) indicates a directional progression from progenitor clusters toward differentiated dI1 neurons along the dI1 arm, with cells at the base showing high outward velocity and low latent time, and those at the tip showing reduced velocity and high latent time. This supports a coherent pseudotemporal ordering of dI1 cells at this single developmental stage and provides a framework to interrogate stage-associated transcriptional programs.

#### Transcriptional dynamics along the dI1 developmental trajectory correlate with spatial cell localization

To track gene expression changes along the dI1 trajectory, pseudotime was assigned to each cell using Slingshot (Fig. 2A). Next, a subset of 954 genes showing significant variation along the control trajectory was identified (Fig. 2B) and assembled into eight groups based on their peak expression along the pseudotime axis using K-means clustering. Whereas in the developing NT, the earliest progenitors span the apico-basal extent of the epithelium, by E4 progenitors localized to the medial (ventricular) zone, whereas differentiating neurons were found in the lateral domain (Fig. 2C). A composite visualization (Fig. 2D) exemplifies the correlation between protein expression and spatial cell localization highlighting medial (PAX7), intermediate (BRN3A), and lateral (BARHL1) expression profiles. To note is that although BARHL1 immunoreactivity was restricted to laterally located neurons, RNA expression was already weakly detectable in progenitors (Fig. 1D, 2B and Sup. Fig. 2). In situ hybridization (ISH) of additional genes from each expression group (Fig. 2B,E), showed a clear medial-to-lateral shift in signal across the dorsal NT that paralleled a bottom-to-top distribution pattern along the UMAP (Fig. 2E, arrows). These results validate the scRNA-seq dataset and its alignment with known patterns of medio-lateral cellular migration in the early developing spinal cord ^39^.

**Figure 2.**
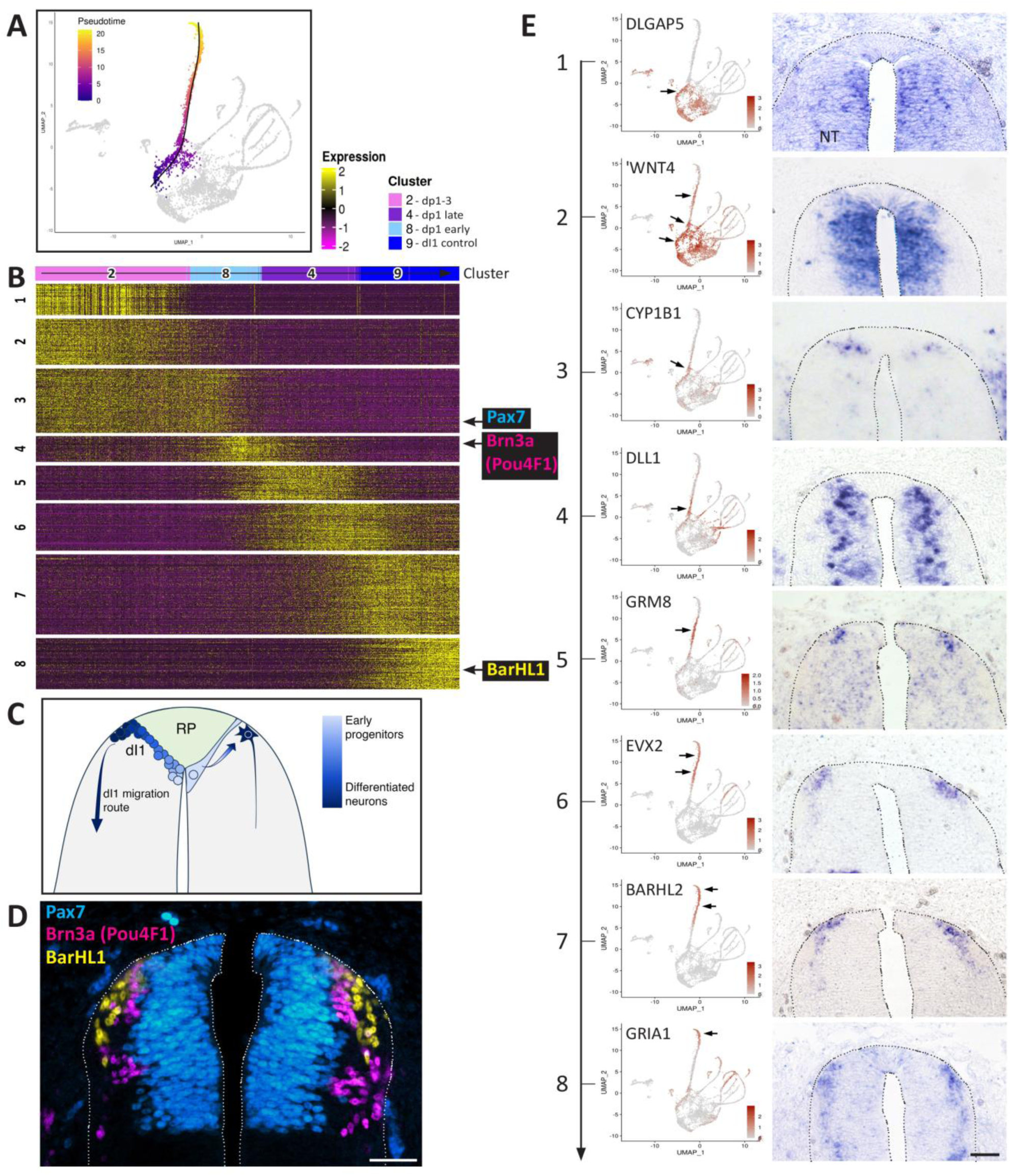
Temporal gene expression patterns during dI1 neuronal differentiation reveal molecular shifts that correlate with spatially organized cellular positioning. A) UMAP of the control cells. Trajectory inference with cluster 2 set as the starting cluster and cluster 9 set as the end cluster. The fitted principal curve is shown in black. Cells are colored according to pseudotime. B) Expression of genes that change along the dI1 trajectory. Cells are ordered by pseudotime and cluster number is indicated as top annotation. Arrows point to the positions of genes shown in C. C) Schematic of dI1 interneuron development ventral to the roof plate (RP). The early stage (right) shows pseudostratified progenitors spanning the apico–basal axis and a basally positioned prospective neuron. The advanced stage (left) illustrates dI1 progression from medial progenitors through intermediate neuroblasts to differentiated lateral neurons, followed by ventral migration (arrow). D) Composite image illustrating the sequential expression of Pax7 (blue), Brn3a (magenta), and BarHL1 (yellow) in distinct domains. E) ISH of the 8 distinct gene groups identified in B. For each cluster, one representative gene is shown, with corresponding expression levels projected onto the UMAP. Arrows in UMAPs indicate expression sites within dI1 interneuron clusters. In all cases there is a correspondence between expression in UMAPs and in the embryo. Abbreviations, NT, neural tube. Scale bar, 50 μm.

#### Stage-specific functional pathways in the developmental trajectory of dI1 interneurons

To identify key biological processes during dI1 interneuron maturation, we performed pathway enrichment analysis on stage-specific gene expression patterns (Fig. 2B) using Enrichr. This approach enabled us to map critical functional programs to successive developmental phases.

Early phases were enriched with genes related to mitosis and DNA replication and repair, consistent with the proliferative nature of neural progenitors (Sup. Fig. 5A-D). Furthermore, a gene subset associated with the mitotic pathway was evident in more advanced clusters; these genes, such as *BTG2, GADD45G, LZTS1* or *DPF3* (Sup. Fig. 5A) are initiated and/or participate in cell cycle arrest and in the switch from proliferation to neurogenesis ^40–42^. These phases were also characterized by high expression of transcriptional regulators and repressors of neuronal differentiation programs such as *SOX2, SOX9*, *PAX7, ZIC1, NOTCH1, HES5, YAP1*, etc. (Sup. Fig. 6A,B). ISH confirmed their localization to ventricular and/or intermediate regions of the NT (Sup. Fig. 6E-R). These and additional genes, establish a molecular environment that maintains the progenitor state while preventing premature neurogenesis.

BMP signaling is well-established as crucial for patterning dI1 interneurons. Studies in avians showed that even late inhibition of BMP activity between E3 and E4 reduces the dp1 population by 60% ^11^. However, the precise temporal requirement for BMP signaling remained poorly characterized. We identified some BMP members as well as BMP signaling targets as significantly enriched in early phases of dI1 development, including *BMP7, GDF11, MSX1/2* and *ID1,3,4* prominently expressed at the beginning of the trajectory (Fig. 3A-F). In the transition to early neurogenesis, BMP inhibitors were upregulated, such as *CHRDL1* ^43^, *DLX1* ^44^, *HIPK2* that represses SMAD1-induced BMP signaling ^45^, and *FSTL4/5* ^46^. Additionally, the BMP and RA-dependent gene *BRINP2*, that negatively regulates mitosis and stimulates neurogenesis ^47,48^ was expressed in late dp1 and in neurons (Fig. 3A,G,H). Thus, whereas our data confirm that BMPs play a critical role during early stages of dI1 development ^49,50^, they further suggest that BMP signaling is dynamically modulated by ligand-repressor interactions during differentiation (Fig. 3A,B and see Discussion). Confirming this notion, pSmad1/5/9, a readout of BMP signaling, was restricted to the ventricular domain of the dorsal NT. Importantly, pSmad1/5/9 showed no overlap with Brn3a (POU4F1) expression, which marks intermediate-stage dI1 cells (Fig. 3I-K). These findings provide evidence for the stage-specific action of BMP signaling in early dI1 development, prior to the onset of neuronal differentiation.

**Figure 3.**
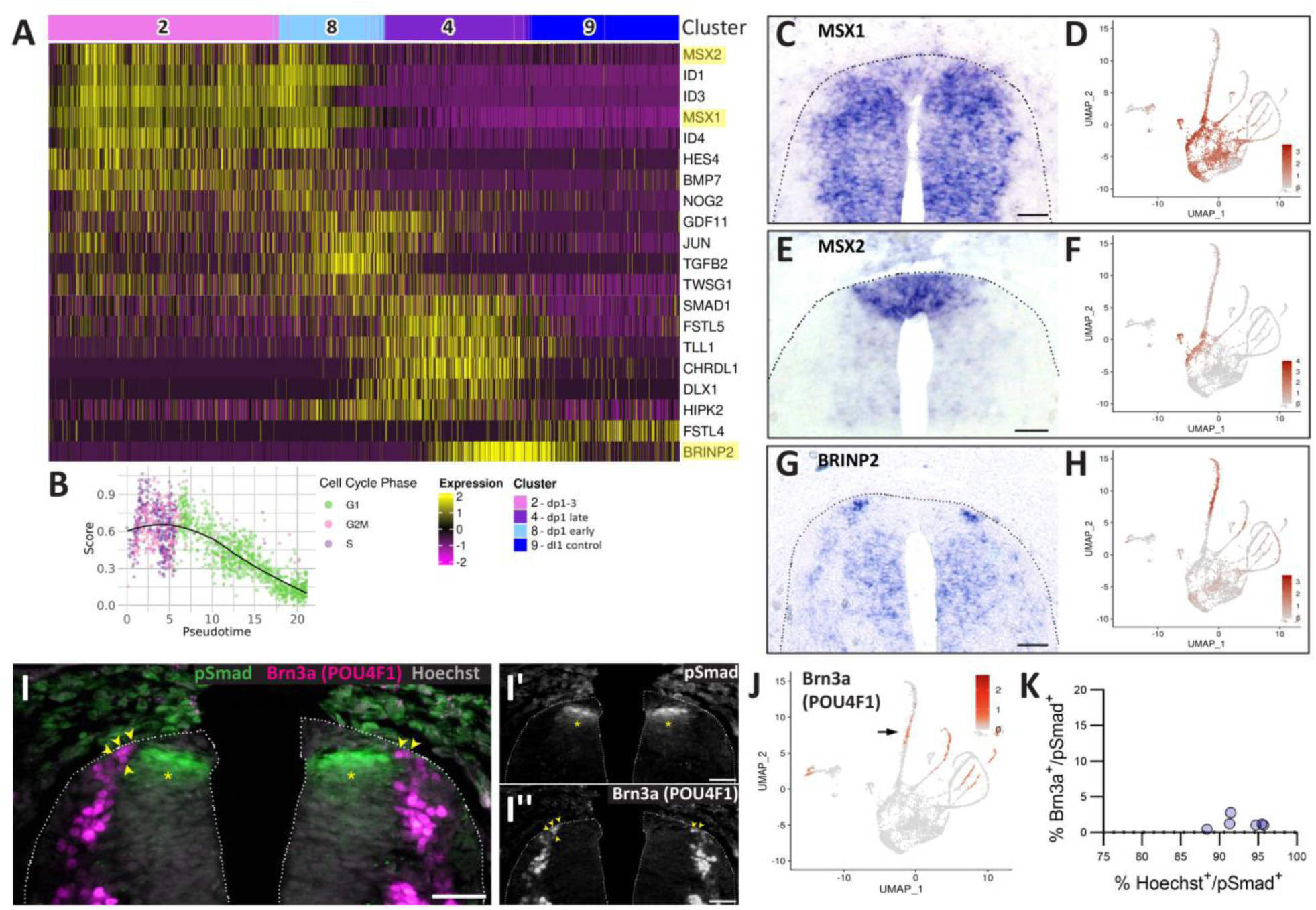
BMP activity is confined to the initial stages of dI1 interneuron development. A) BMP pathway gene expression along the dI1 differentiation trajectory (Seurat-scaled values), with validated genes (C–H) highlighted in yellow. BMP ligands (*BMP7, GDF11, TGFB2*) and direct targets (*MSX1/2, ID1/3/4*) are enriched in progenitor clusters (2, 8), whereas differentiating and differentiated neuron clusters (4, 9) predominantly express BMP inhibitors (*FSTL4/5, CHRDL1, HIPK2, BRINP2*). Cells are ordered by pseudotime, with cluster numbers shown above. B) Average gene expression levels of BMP signaling-related genes along the pseudotime, with cells color-coded by cell cycle phase. C-H) ISH for representative genes of the BMP pathway, with corresponding UMAP projections. I) Immunostaining for pSmad1/5/9 (green, asterisks) and Brn3a (magenta, arrowheads) demonstrates that the two signals are mutually exclusive. Panels I’ and I” represent separate images for the above. J) Expression levels of Brn3a (POU4F1) projected onto the UMAP, arrow indicates dI1 interneurons. K) Quantitative assessment of the spatial overlap between Brn3a^+^ and pSmad^+^ regions, with Hoechst^+^/pSmad^+^ overlap serving as a positive control. N = 6 embryos (7 sections/embryo). Scale bar, 50 μm.

Other central signaling pathways in development exhibit more complex expression patterns. Genes associated with Wnt signaling are expressed in a nuanced trajectory during dI1 development, with a mild downregulation observed across the developmental continuum. Dp1-3 and early dp1 progenitors express *WNT1, WNT4, AXIN2,* and *NDP* (Sup. Fig. 7A,C,E-H), consistent with a role of Wnt ligands in dI1 development ^33^ and with multiple reports positioning their activity downstream of BMP ^6,30,31,51^. Wnt signaling may contribute to later stages of dI1 development when BMP activity is no longer apparent (Fig. 3). For instance, *WNT16* is primarily expressed in cluster 4 (late dp1) with some expression in early dp1 progenitors (Sup. Fig. 7A, I-J), and has been associated with canonical Wnt signaling during cell differentiation ^52^. Late-stage transcription of *RSPO2* and *DACT1* may further modulate Wnt function/s during neuronal differentiation ^53,54^.

Notch pathway genes are enriched in the early trajectory (Sup. Fig. 7B,D). *NOTCH1*, *HES5*, *HES6* and *HEYL* are expressed in dp1-3 and early dp1 clusters. *NOTCH1* is excluded from the RP (Sup. Fig.7B,D,K,L), similar to its ligands *DLL1* and *JAG2,* which are upregulated at intermediate stages across all dorsal interneuron subtypes (Fig. 2D, Sup. Fig 7M-N), marking their neuronal specification. These data in the quail are consistent with known regulatory functions of Notch signaling in maintaining neural progenitor stemness in the mammalian brain ^55^, with its requirement for RP and dI1 development in avians and mice ^37^, and in establishing the RP-dI1 boundary in the spinal cord ^9^.

#### The dynamics of pathways associated with neuronal development

Neuronal development involves dramatic transformations in cell morphology and positioning within the neural tissue. As neuroepithelial progenitors differentiate into dorsal interneurons, they undergo a coordinated sequence of cellular remodeling events. First, progenitors migrate laterally from the ventricular zone, progressively losing their epithelial characteristics. Subsequently, as these cells acquire neuronal identity, they migrate ventrally to their final position ^16^. Our analysis showed overall upregulation of genes involved in cytoskeletal dynamics, extracellular matrix (ECM), and cell migration along the dI1 trajectory (Sup. Fig. 8). These genes exhibited distinct temporal expression, indicating multiple waves of cellular reorganization during dI1 development. An early wave of cytoskeleton-associated genes corresponded to cell division, actin remodeling, cytoneme formation, and motility *(NHS, HMMR, DIAPH3, MYO10, CYR1A)*, followed by a later wave linked to neuronal microtubule assembly, axonal growth, and vesicle transport *(NEFM, KIF26B, MID1, STMN3, MAPRE2, PPFIA2, NAV1*, etc.) (Sup. Fig. 8A,D,G,H). Likewise, cell adhesion and ECM genes associated with growth and proliferation (e.g., *PCDH19, CCN2, FAT1*) were predominantly expressed in dp1–3 and early dp1 clusters, whereas neurite growth–related genes (e.g., *LAMC2, HS6ST2, NFASC, THSD1*) emerged during later neurogenesis (Sup. Fig. 8B,E). Distinct subsets of “migratory” genes, including ECM, cytoskeletal, and guidance molecules, displayed cluster-specific distributions across dI1 populations (Sup. Fig. 8C,F), indicating that cell migration occurs throughout all stages of dI1 ontogeny.

#### Mechanisms of cell survival in neuronal development

Cell survival pathways tightly regulate progenitor and neuronal population size during development ^56^. In the dI1 lineage, genes regulating apoptosis peak at the intermediate (clusters 8 and 4) stages (Fig. 4A,B). Although apoptosis is classically associated with later elimination of excess post-mitotic neurons ^57^, its early role is unclear. Notably, the stages show concurrent upregulation of pro-apoptotic genes (e.g., *JUN, TGFB2, PLK3*) and anti-apoptotic regulators [*BCL2L1 and BCL2L10* ^58^*, ITGA6* ^59^*, INPP4A* ^60^, and *MEF2C* ^61^ (Fig. 4A)]. Yet, cleaved caspase-3 staining revealed no apoptotic cells in the NT at E4 or E4.5, despite distinct apoptosis in adjacent dorsal root ganglia (Fig. 4C–E). Thus, the apoptotic program appears transcriptionally primed but inactive during specification, which may allow later activation or relate to proposed non-apoptotic functions in neuronal differentiation (see Discussion).

**Figure 4.**
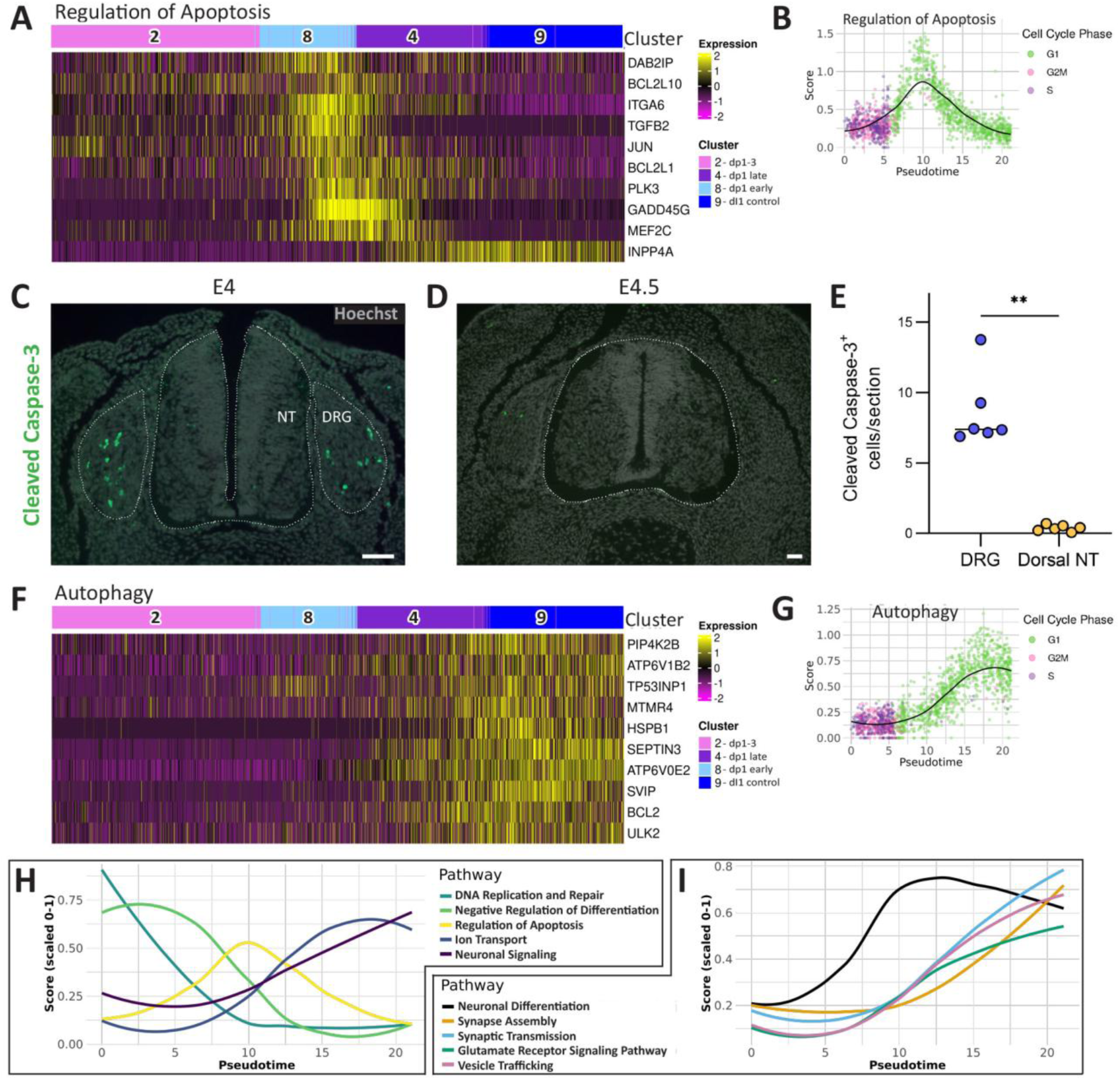
Control of cell survival/apoptosis during dI1 interneuron development. A) Heatmap of genes associated with the regulation of apoptosis (Seurat scaled expression values). Cells are ordered by the pseudotime and cluster number is shown as top annotation. B) Average gene expression levels of “Regulation of Apoptosis” pathway genes, with cells color-coded by cell cycle phase. C-E) Immunostaining for Cleaved Caspase3 (green) at E4 (C) and E4.5 (D). Positive cells are observed in the DRG but not in the NT. E) Quantification of Cleaved Caspase3^+^ cells at E4 in NT and DRG. Mann-Whitney test; **p < 0.01. N = 6 embryos/ group, 14-20 sections/ embryo. F) Heatmap of genes associated with the autophagy. Most genes are upregulated in differentiating and differentiated neurons. G) Average gene expression levels of autophagy pathway genes, cells color-coded by cell cycle phase. H-I) Comparative analysis of expression dynamics for key regulatory pathways controlling dI1 differentiation. Plotted are pathway average scores along the pseudotime. Abbreviations, DRG, dorsal root ganglion, NT, neural tube. Scale bar, 50 μm.

Autophagy supports cell survival by recycling proteins and organelles ^62^. Autophagy-related genes were significantly enriched in late dp1 cells and in dI1 neurons (Fig. 4F–G), consistent with reports that autophagy is required for differentiation-associated cellular remodeling ^63^. Further experiments will be required to clarify the mechanisms underlying the dynamics of both apoptotic and autophagy genes described here.

#### Neuronal functions induced during differentiation

In the E4 embryo, expression of genes involved in neuronal specification and differentiation follows a clear temporal progression with three distinct yet partially overlapping phases (Sup. Fig. 9A,D). Early patterning genes, including *PAX7* and *OLIG3*, establish dorsal neural identity. This is followed by intermediate-stage specification factors such as *INSM1* ^64^*, ATOH1, NEUROG2, BRN3, CBL1* and *ZEB1*, which direct cells toward a neuronal fate. Then, differentiation genes including *LHX9, BARHL1/2, NGFR* and *PLK2* become upregulated ^5,6,15,19,65,66^. Notably, *LHX2* is highest in the transition from progenitors to differentiation (cluster 4) but downregulated in the terminally differentiated neurons (cluster 9) (Sup. Fig. 9A, O,P) ^16^.

A clear temporal shift in gene-expression specificity is observed along the differentiation timeline. Early patterning genes are broadly expressed across multiple neuronal subsets including the relatively restricted factor *OLIG3* that spans progenitor domains dp1–3. In contrast, genes conferring precise neuronal subtype identity emerge primarily at intermediate to late stages (clusters 8 and 4).

Genes associated with mature neuronal functions are activated predominantly during later development (clusters 4 and 9). Pathways involved in neuronal signaling, glutamate receptor signaling, Semaphorin–Plexin signaling, neuronal projections, ion transport, and synaptogenesis are coordinately upregulated toward the end of the differentiation trajectory (Fig. 4H–I, Sup. Figs. 9–11). The mechanisms underlying this synchronized activation remain unclear, although engagement of the neuronal differentiation pathway immediately beforehand (Fig. 4I) may coordinate this transition.

#### Comparative transcriptional analyses reveal that dI1, dI2, and dI4 interneuron subtypes exhibit both shared and distinct regulatory programs

Next, we asked whether the transcriptional patterns seen in dI1 interneurons are exclusive or common to other neuronal types by comparing them with dI2 and dI4, two populations with continuous pseudotime trajectories (Sup. Fig. 12). Genes regulating apoptosis and autophagy showed similar expression dynamics across all three subtypes, indicating common survival mechanisms, though some regulators differed: *BCL2L10* was more pronounced in dI4, while *ITGA6* and *MEF2C* were primarily expressed in dI1 (Sup. Fig. 12A–B, E–F).

BMP signaling is required for the development of dI1-dI3 but not for dI4-6 interneurons ^19,21,28,31^. Consistently, BMP-related gene expression matched closely between dI1 and dI2 but differed from dI4. For example, BMP inhibitors such as *NOG2* and *FSTL4* were detected in emerging dI1 and dI2 neurons, and *HIPK2, FSTL5, TLL1, CHRDL1,* and *DLX1* mostly in dI1, marking BMP signaling cessation. Early-expressed genes including *ID1, ID3, MSX1, HES4*, and *BMP7* showed similar but lower expression in early dI4 progenitors, suggesting residual BMP influence in early dI4 development despite later independence (Sup. Fig. 12C,G).

Genes involved in neuronal specification and differentiation generally displayed overlapping expression among dI1, dI2, and dI4, though a distinct subset, such as *INSM2, WNT16, DLX1*, *BRN3*, *LHX2*, *LHX9*, *BARHL1*, *BARHL2*, *PLK2* and *ATOH1,* specifically defined dI1 identity (Sup. Fig. 12D,H). DNA replication and repair genes were upregulated early in all trajectories, while neuronal signaling genes were consistently higher in later maturation stages, reflecting shared patterns of proliferation and specialization at the extremities of the trajectory (Sup. Fig. 12I-J).

### RA exerts stage-dependent effects on the dynamics of dI1 interneuron development

#### Comparative analysis of control versus RARα403-treated conditions

RA signaling plays a central role in neural development ^67,68^ and was recently implicated in cellular fate segregation within the dorsal NT ^9^. Our scRNA-seq analysis encompassed 10 control embryos as well as 10 embryos subjected to RA inhibition through electroporation of a dominant-negative truncated human RARα receptor (RARα403). This construct inhibited by 88% the intensity of RA activity compared to controls ^26^.

Control and RARα403 samples clustered similarly at most developmental stages, except at terminal differentiation, where control (cluster 9) and RARα403 (cluster 15) dI1 populations separated distinctly (Sup. Fig. 13A-B). Similar to the cell cycle behavior monitored under control conditions, most mitotic progenitors in the combined control and treated samples were located in the main body of the UMAP with only few cycling cells along the treated arm (Sup. Fig. 13C, arrow). Furthermore, RNA velocity and latent time analyses confirmed a similar developmental trajectory in both conditions (Sup. Fig. 13D,E), enabling direct stage-by-stage comparison of gene expression throughout dI1 development.

#### Functional modules along the dI1 developmental trajectory are affected by RA activity

To investigate the role of RA signaling in dI1 interneuron development, we conducted a comprehensive differential expression analysis comparing control and RARα403-treated samples across the entire developmental trajectory. This revealed 571 genes whose expression was significantly altered by perturbation of RA signaling (Sup. Fig. 14). Genes with stage-specific expression patterns along the trajectory are presented in groups 1-12, while those with consistent expression across developmental stages are shown in groups 13-16 (Sup. Fig. 14). Additionally, genes were classified based on their directional response to RA inhibition (either up- or down-regulated). These RA-responsive genes were then systematically grouped into six functional modules to elucidate how RA regulates various aspects of dI1 development (see below and Figs. 5-7, Sup. Fig. 15).

**Figure 5.**
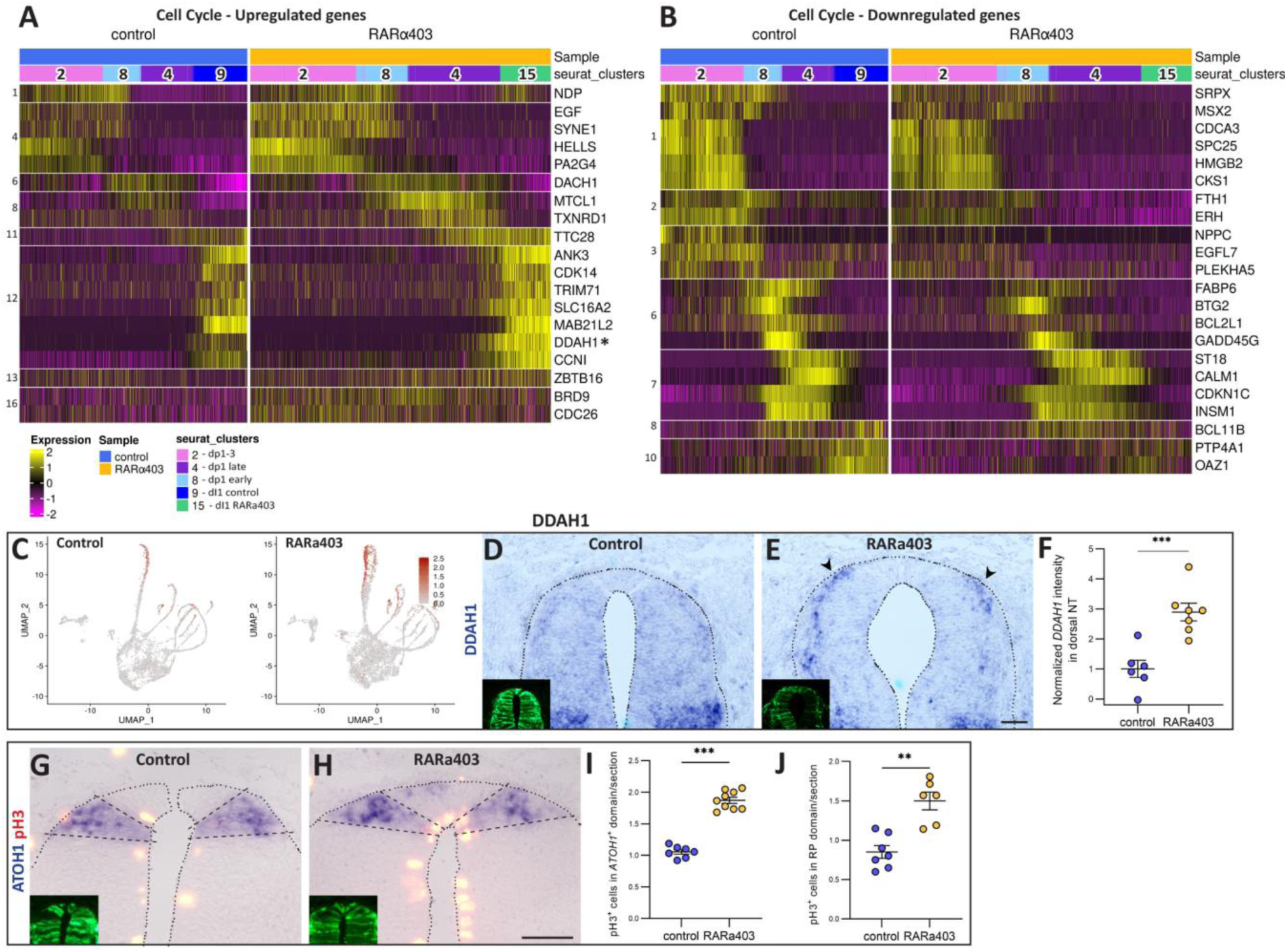
RA affects the expression of cell cycle-related genes along dI1 interneuron development. A-B) Heatmaps of genes associated will Cell Cycle regulation following RA inhibition. Shown are upregulated genes (A) and downregulated genes (B) in RARα403-treated samples compared to control. Genes are classified according to the gene groups defined in Sup. Figure 14. C-F) Validation of *DDAH1* expression changes (marked with asterisk in A). UMAP projection of *DDAH1* expression levels from single-cell transcriptome data (C). For spatial validation, embryos were electroporated with control PCAGG or RARα403 at E2.5, fixed at E4, and analyzed by ISH for *DDAH1*. Control embryos show minimal *DDAH1* expression in the dorsal NT (D), while RARα403-treated embryos exhibit expression in the differentiating dI1 domain (E, arrowheads), consistent with the bioinformatic analysis (C). Insets in D and E illustrate the corresponding electroporated NTs. Quantitative analysis of *DDAH1*^+^ staining intensity in dorsal NT is shown in (F). G-I) Cell proliferation (pH3^+^ mitotic nuclei) in the *ATOH1*^+^ domain of control (G) and RARα403-treated (H) embryos. Insets show GFP-electroporated cells in corresponding sections. I) Quantification showing a significant upregulation of mitoses in dI1 interneurons. J) Quantification of cell proliferation in the RP domain, dorsal to the *ATOH1*^+^ area. Student’s t-test was applied; ***p < 0.001. N = 11-15 sections per embryo in 6-11 embryos were analyzed in the different groups. Scale bar, 50 μm.

#### Cell cycle regulation

RA inhibition led to stage-dependent changes in cell cycle regulation (Fig. 5A,B). In dp1-3 and early dp1 (clusters 2 and 8), both up- and downregulated genes consisted of positive modulators of cell cycle progression, with upregulated genes including *NDP, EGF, HELLS,* and *SYNE1*, and downregulated genes such as *CDCA3, SPC25, HMGB2, CKS1, FTH1, ERH, EGFL7,* and *PLEKHA5*. At intermediate stages (dp1 early and dp1 late, clusters 8 and 4), downregulated genes were mostly negative regulators of the cell cycle (*BTG2, GADD45G, CDKN1C, INSM1, BCL11B*). During differentiation, RARα403 treatment primarily upregulated genes stimulating cell cycle activity, including *CDK14, ANK3,* and *DDAH1* (Fig. 5A,C-F). To directly assess the effect of RARα403 on cell proliferation, we quantified the number of pH3-positive mitotic nuclei within the dI1 population, identified by *ATOH1* expression. RARα403 significantly increased the number of mitotic cells in the *ATOH1*-positive domain, corroborating the bioinformatic findings (Fig. 5G–I). In addition, increased proliferation was also observed in the RP (Fig. 5J), further supporting our previous results ^26^. Together, these findings suggest that RA modulates cell-cycle dynamics throughout dI1 development, potentially influencing the timing of cell-cycle exit.

#### RA Coordinates Multiple Signaling Pathways During dI1 Development

Genes impacted by RA inhibition and involved in cell signaling were mainly part of the BMP, Wnt, Notch, and RA pathways (Fig. 6). Notch activity was enhanced in the absence of RA activity, as evidenced by the upregulation of *NOTCH1* and *JAG1* in early clusters 2 and 8 and the Notch receptor *MINAR1* in the late progenitors and in neurons (clusters 4 and 15). This pattern suggests that RA continuously suppresses Notch signaling in dI1 progenitors, consistent with recent findings showing that RA restricts Notch activity to the boundary between RP and dI1 interneurons ^9^.

**Figure 6.**
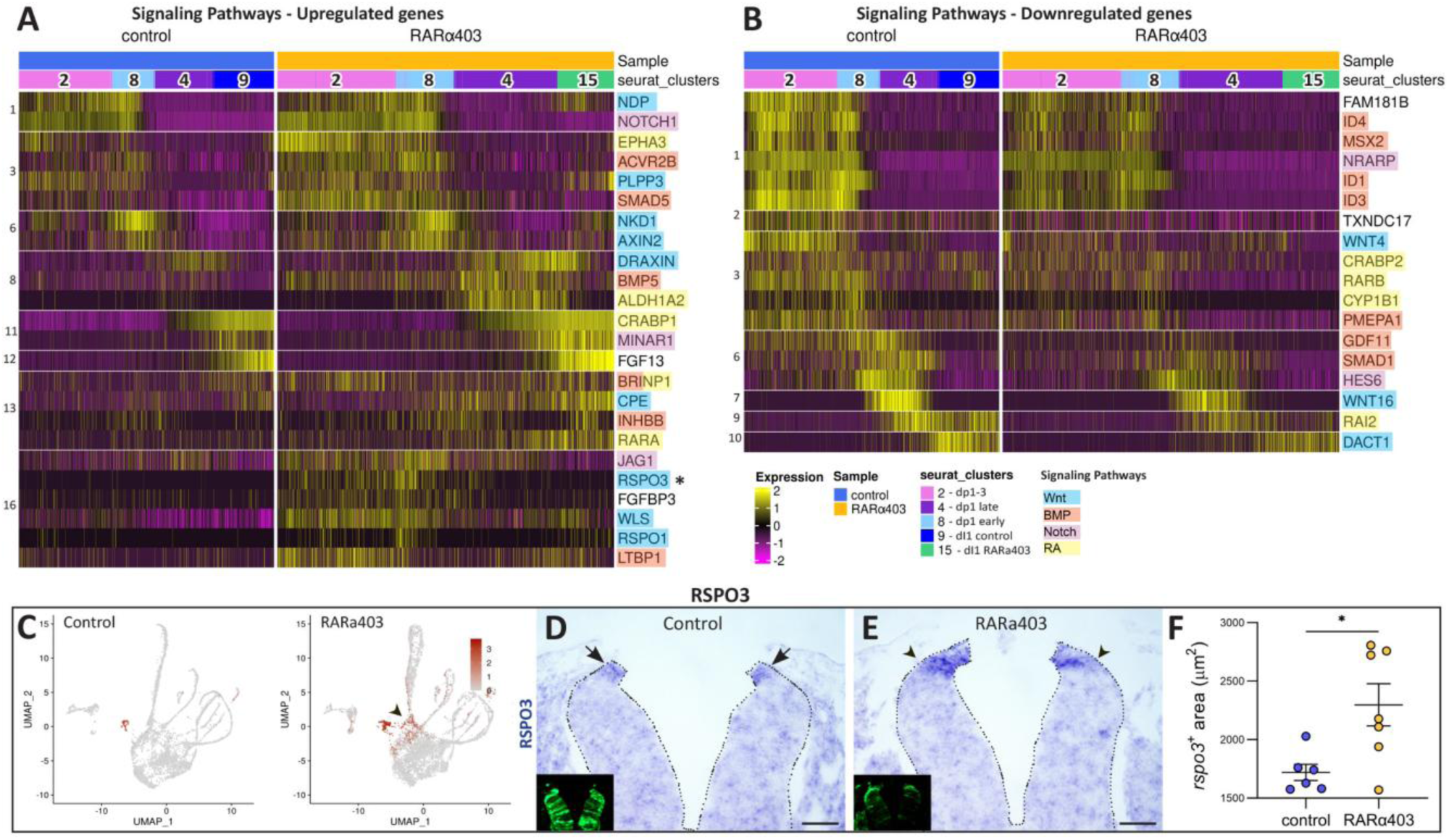
RA inhibition alters expression of key developmental signaling pathways in dI1 interneurons. A-B) Differential gene expression of signaling pathways following RA inhibition, showing upregulated (A) and downregulated genes (B) in RARα403-treated samples compared to control. Genes are classified according to the groups defined in Sup. Figure 14. Genes are color-coded by main pathways. C-F) Validation of RSPO3 expression changes (asterisk in A). UMAP projection of *RSPO3* expression levels from single-cell transcriptome data (C). For spatial validation, embryos were electroporated with control PCAGG or RARα403 at E2.5, fixed at E4, and analyzed by ISH for *RSPO3*. Control embryos show *RSPO3* expression in the RP (D, arrows), while RARα403-treated embryos exhibit expanded expression in the progenitor dI1 domain (E, arrowheads), consistent with the bioinformatic analysis (C). Insets in D and E illustrate electroporated NTs. Quantitative analysis of *RSPO3*^+^ shown in (F). Welsh’s t-test; *p < 0.05. N = 6,7 embryos for control and treated groups, respectively. 15-26 sections/embryo. Scale bar, 50 μm.

The effects of RA inhibition on the RA pathway were complex. In the early half of the trajectory, the binding protein *CRABP2*, the receptor *RARB*, and the RA-producing enzyme *CYP1B1* were all downregulated. In contrast, at later stages, RA inhibition led to increased expression of the RA-producing enzyme *ALDH1A2* in late cluster 4, and upregulation of the binding protein *CRABP1* and receptor *RARA* in differentiating neurons. These results suggest that RA boosts the expression of its own positive regulators during early development, but reduces them at later stages, indicating a greater need for RA activity in early progenitors that decreases as development progresses.

RA inhibition caused the upregulation of both positive Wnt effectors (*NDP, AXIN2, PLPP3*) and negative regulators (*DRAXIN, CPE, NKD1*), alongside a downregulation of the inhibitor *DACT1* in dI1 neurons. Dp1-3 and early dp1 clusters showed increased expression of the positive regulators *RSPO1* and *RSPO3* (Fig. 6A,C-F), while *WLS*, involved in Wnt ligand secretion, was upregulated throughout the developmental trajectory. Collectively, these findings suggest that RA suppresses Wnt activity in dI1 interneurons. This could reflect an RA-mediated brake on Wnt-mediated proliferative signaling, as previously shown for the RP ^26^, thus coordinating the transition from proliferative progenitor states to post-mitotic neuronal differentiation.

BMP pathway components were predominantly downregulated by RARα403 treatment, including *ID1, ID3, ID4, MSX2, GDF11, SMAD1,* and *PMEPA1*. Notably, recent findings suggest a role for the positive Wnt effector RSPO3 in inhibition of BMP signaling. This is in line with the observed upregulation of *RSPO3* in dp1-3 and early dp1 clusters and the parallel downregulation of BMP targets (Fig. 6) ^69^. Therefore, RA upregulates BMP-related genes in dI1 progenitors, suggesting that BMPs promote early cell cycle progression and specification.

*BRINP1*, a gene induced by RA and BMP signaling, was upregulated after RA inhibition, especially in dp1-3 (Fig. 6A). As BRINP1 suppresses cell cycle progression and neurogenesis ^70^, these findings are consistent with a potential influence of RA on progenitor proliferation and neuronal differentiation through modulation of *BRINP1* expression.

#### Neuronal Specification, Differentiation and Signaling

RA inhibition led to upregulation of genes involved in neuronal differentiation and signaling primarily in clusters 4 and 15 including ion channels (*KCND2, NKAIN3, KCNH6, SCN2A, SCN3A, CACNA2D1*), synaptic markers (*RPH3A, CHGB, CBLN2, NXPH2, UNC13C, SV2C),* and neurotransmitter receptors (*GRIN2A, GABRA3, GABRE, GRIA1*) (Sup. Fig. 15A-M). In contrast, RA inhibition did not significantly alter *ATOH1* mRNA expression (Sup. Fig. 15N-P), suggesting that RA may not substantially affect dI1 specification, but may instead act later to negatively modulate aspects of terminal differentiation.

#### Temporal control of cytoskeletal architecture, extracellular matrix and cell adhesion

RA exerts stage-specific control over genes governing cellular architecture and extracellular interactions. The majority of cytoskeleton and ECM-related genes affected by RA perturbation were upregulated following RA inhibition, with this effect most pronounced at advanced developmental stages (Fig. 7A-F). Key upregulated genes included the microtubule-associated protein *DCX*, tubulin isoforms, laminin family members, the synaptic organizer *AGRN*, *MYH15* and the cytoskeletal regulator *FIGN*. This pattern indicates that endogenous RA activity serves as a brake on cytoskeletal reorganization programs, particularly as neurons approach terminal differentiation. This regulatory mechanism may be associated with RA negatively affecting terminal neuronal differentiation (Sup. Fig 15) and be essential for proper timing of morphological maturation, ensuring that cytoskeletal remodeling occurs in coordination with differentiation.

**Figure 7.**
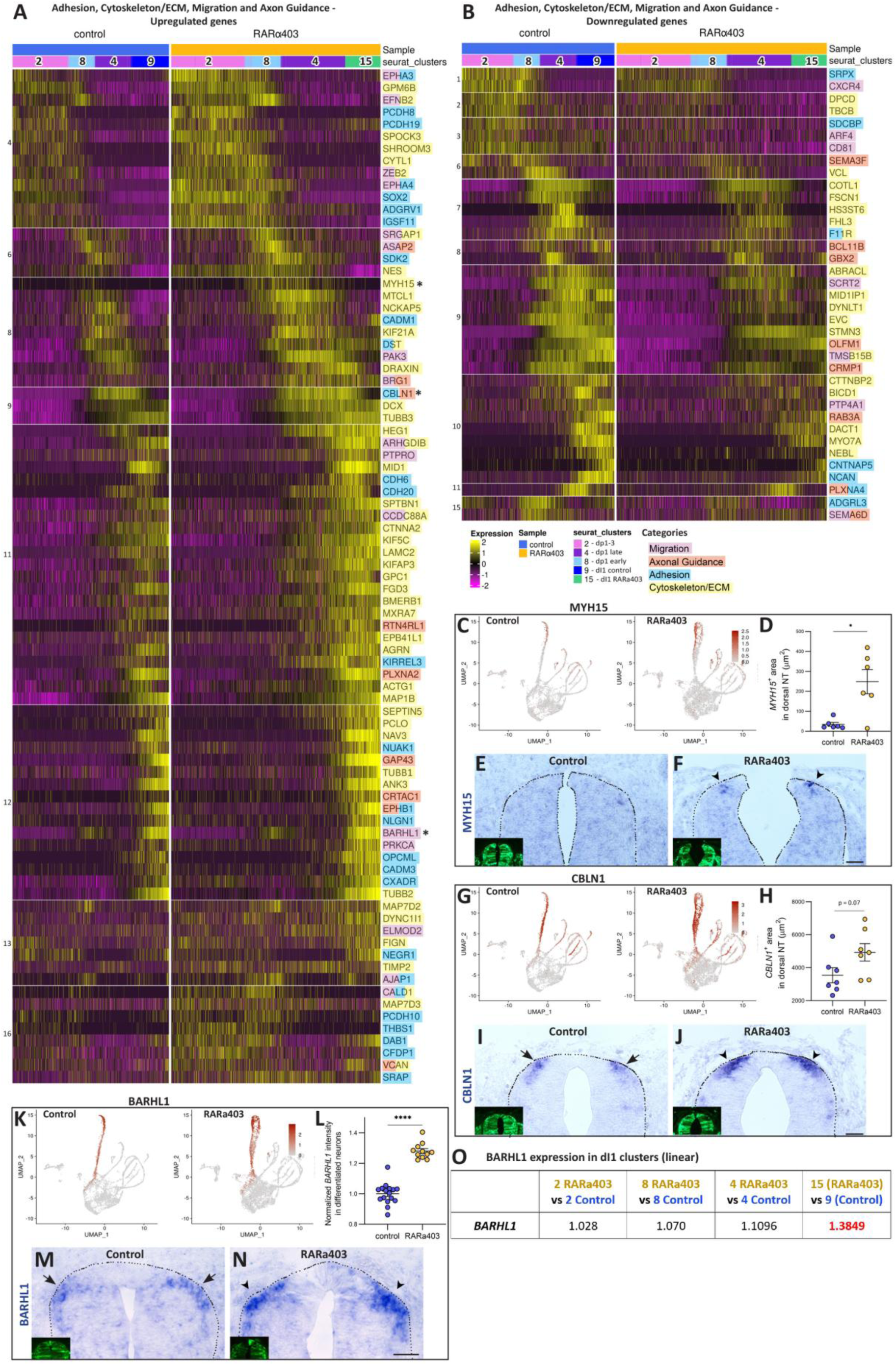
Dynamic Regulation of Structural and Guidance Mechanisms upon inhibition of RA signaling. A-B) Heatmaps of genes affected by RA inhibition and associated with Adhesion, Cytoskeleton, Extracellular Matrix, Migration and Axon Guidance. Shown are upregulated genes (A) and downregulated genes (B) in RARα403-treated samples compared to control. Genes are classified according to the gene groups defined in Sup. Figure 14 and are color-coded by main pathways. C-N) Validation of *MYH15* (C-F), *CBLN1* (G-J) and *BARHL1* (K-O) expression changes (genes marked with asterisk in A). Embryos were electroporated with control PCAGG or RARα403 at E2.5, fixed at E4, and analyzed by ISH. RARα403-treated embryos exhibit enhanced and expanded expression in the differentiating dI1 domain (F,J,N, arrowheads), consistent with the bioinformatic analysis (C,G,K). Insets in E,F,M and I,J,N illustrate the corresponding electroporated NTs. Quantitative analysis of *MYH15*^+^, *CBLN1*^+^, and *BARHL1+* area is shown in D, H and L. O) Fold-increase in *BARHL1* mRNA expression in each cluster. Welsh’s t-test was applied; *p < 0.05, ****p<0.0001. N = 6,6 (*MYH15*), 7,7 (*CBLN1*) and 16,11 (*BARHL1*) embryos for control and treated groups. 8-17 (*MYH15*) and 10-20 (*CBLN1* and *BARHL1*) sections per embryo were analyzed. Scale bar, 50 μm.

In contrast to the late-stage effects on cytoskeletal genes, the influence of RA on cell adhesion molecules was most prominent during early developmental stages. Cell adhesion-related genes such as *PCDH8, PCDH19, ADGRV1, IGSF11*, were predominantly upregulated following RA inhibition in dp1-3 and early dI1 clusters, and so were *EPHA3* and *EPHA4* that mediate adhesion and axon guidance (Fig. 7A), suggesting that endogenous RA activity normally suppresses adhesive interactions during the initial phases of dI1 development. This effect may facilitate the dynamic cellular rearrangements necessary for proper progenitor positioning and initial specification events.

#### Complex Regulation of Axonal Guidance Networks

RA control over axonal guidance machinery revealed complex regulatory patterns (Fig. 7A,B). The Semaphorin-Plexin pathway, which mediates axonal repulsion, was generally downregulated following RA inhibition, suggesting that RA normally promotes expression of these repulsive guidance cues. Members of the Eph/ephrin family (*EPHA3, EPHA4, EFNB2*), which also function as axon repellents, showed the opposite pattern, being upregulated in early dp1-3 and dp1 clusters following RA inhibition. So was *CBLN1*, that acts as an axonal growth and guidance cue in early neural development (Fig. 7A,G-J). Additionally, *BARHL1*, involved in conferring commissural identity to dorsal interneurons ^66^, was significantly upregulated in differentiated neurons of cluster 15 (Fig. 7K-O).

#### RA activity is necessary for proper ventral migration of dI1 interneurons

The significant effects of RA on cytoskeletal, adhesion, and migration-related gene modules prompted us to investigate whether these molecular changes translate into altered cellular behaviours. We focused on the ventral migration of dI1 interneurons. Born as the dorsal-most spinal neuron subtype, dI1 interneurons migrate ventrally to establish their final residence in the deep dorsal horn. This migration begins at embryonic day 4 (E4) of avian development, continues from E4.5 to E7, and concludes by E8 ^16^.

To assess the role of RA in dI1 migration, we first mapped BarHL2^+^ dI1 cells at E5 at the flank level. RA inhibition resulted in a marked accumulation of dI1 cells at their origin in the dorsal NT compared with control neurons that efficiently migrated ventralward (Fig. 8A-F). To distinguish a failure to migrate from a delay, we extended our analysis to E6. In RARα403-treated embryos, fewer dI1 cells were positioned within the ventral high-density region typical of controls (Fig. 8G-L). Some cells were still migrating, as evidenced by their elongated nuclear morphology (Fig. 8M-N’) ^71^, while others had migrated to more ventral positions than observed in controls (Fig. 8G-L), suggesting aberrant migration trajectories.

**Figure 8.**
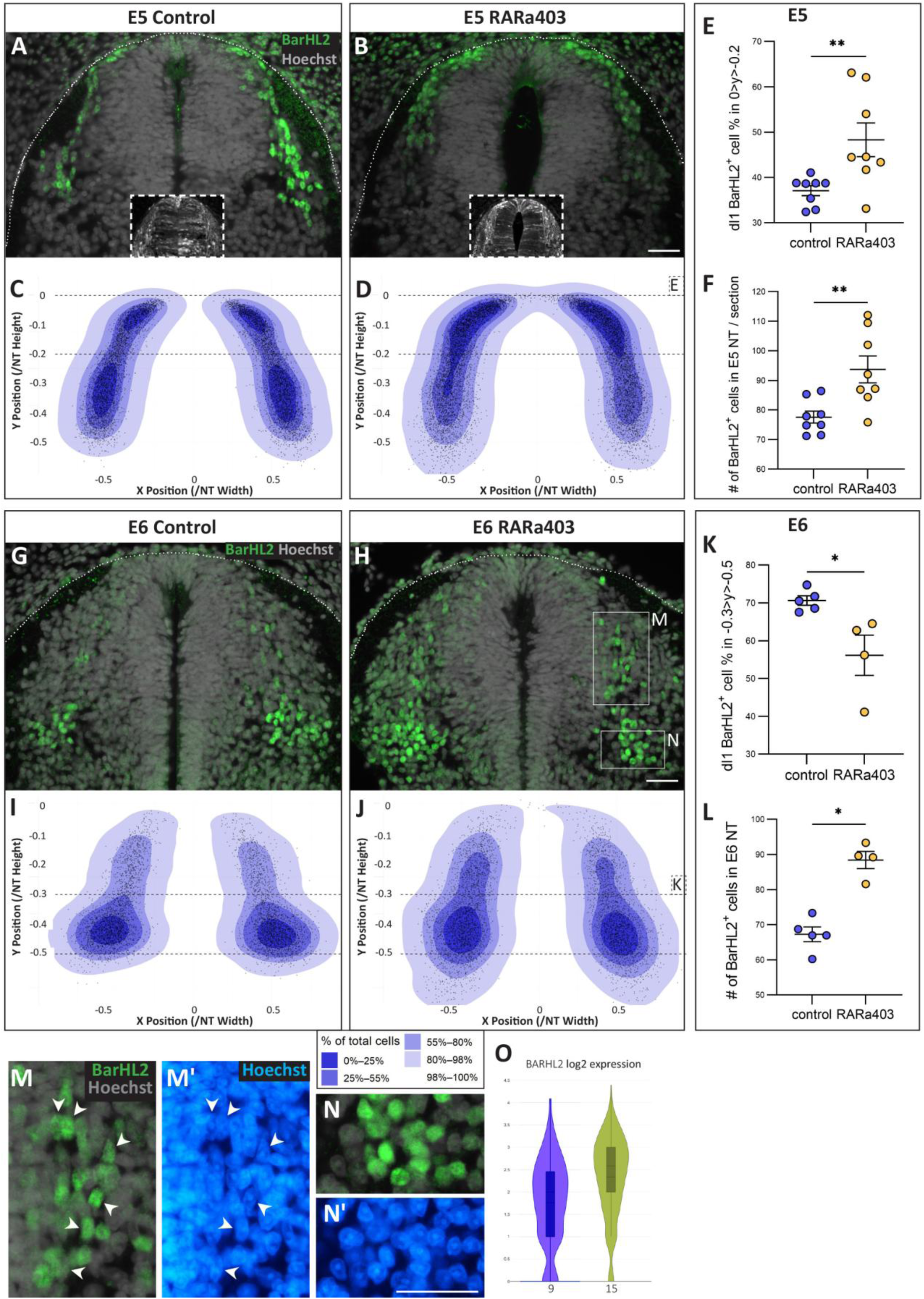
RA signaling is required for proper ventral positioning of dI1 interneurons. A-F) Embryos were electroporated at E2.5 with either control PCAGG (A, C) or RARα403 (B, D), fixed at E5, and immunostained for BarHL2. A, B) Representative spinal cord sections showing the distribution of BarHL2⁺ dI1 interneurons. C, D) Corresponding 2D density maps reveal altered distribution in RARα403-treated embryos. Contours indicate increasing cumulative percentages of the total cell population. Black dots indicate individual cell positions. Dashed lines demarcate the dorsal region quantified in E. N = 7448 cells (control) and 8998 cells (RARα403) from 8 embryos/group; 15 sections/ embryo. E) Proportion of BarHL2⁺ cells retained within the dorsal 20% of the NT. Mann-Whitney test; **p < 0.01. F) Total number of BarHL2^+^ cells at E5. Mann-Whitney test; **p < 0.01. G-L) Embryos were electroporated and processed as above, then fixed at E6. G, H) Representative BarHL2-stained sections from control (G) and RARα403 (H) embryos. I, J) 2D density maps demonstrate persistent distribution abnormalities in RARα403-treated embryos. Contours as in (C, D). Dashed lines indicate the region quantified in (K). N = 5045 cells from 5 embryos (control) and 5306 cells from 4 embryos (RARα403); 15 sections/embryo. (K) Percentage of BarHL2^+^ cells localized to the ventral highest-density zone in control group. Mann-Whitney test; *p < 0.05. (L) Total number of BarHL2^+^ cells at E6. A significant increase is observed in RARα403-treated embryos. Mann-Whitney test; *p < 0.05. (M-N’) High magnification of the boxed regions in (H), showing elongated BarHL2^+^ nuclei, typical of migrating cells (M-M’), compared to the rounded nuclei of settled cells (N-N’). (M’,N’) Hoechst nuclear staining. O) Violin plot depicting the increase in *BARHL2* mRNA expression in differentiated cluster 15 (treated) compared to 9 (control). Scale bar, 50 μm.

Notably, the total number of BARHL2⁺ cells was increased in RARα403-treated embryos at both developmental timepoints (Fig. 8F,L). This increase likely reflects the enhanced proliferation of dI1 progenitors, as indicated by both our transcriptomic analysis and experimental validation (Fig. 5), together with the increased neuronal differentiation observed following RARα403 treatment (Sup. Fig. 15; Fig. 7K–O). Consistent with this interpretation, *BARHL2* mRNA expression was also elevated under these conditions in terminally-differentiated clusters (Fig. 8O).

To assess whether the observed migration effect is specific to the dI1 neuronal subpopulation, we co-stained embryos for Isl1, a marker of differentiated dI3 neurons and motoneurons (Sup. Fig. 16A-B). We found that dI3 neurons – another dorsal subpopulation that typically migrates ventrally, also accumulated near their dorsal origin after RA inhibition (Sup. Fig. 16C-G). However, unlike dI1 cells, the total number of Isl^+^ dI3 neurons and motoneurons decreased in treated embryos (Sup. Fig. 16H,K). This is consistent with the known role of motoneuron-derived RA in promoting their own proliferation and differentiation ^72–75^, and also indicates that RA may differentially regulate cell numbers in dI1 and dI3 populations (see Discussion).

This distinction is further supported by a direct comparison of proliferation and migration defects. The increase in dorsally-retained dI1 cells in RARα403-treated embryos (1.54-fold relative to controls) exceeded the increase in total dI1 cell number (1.2-fold), indicating that increased proliferation alone cannot account for the dorsal accumulation phenotype (Sup. Fig. 16L). Furthermore, total cell number and dorsal retention were positively correlated in dI1 neurons but negatively correlated in dI3 neurons, supporting the conclusion that proliferation and migration are regulated independently in these cell subtypes (Sup. Fig. 16M-N; see Discussion).

Together, these results establish RA as a critical modulator of dI1 interneuron migration, linking molecular changes identified in proliferation, cytoskeletal, migratory and adhesion traits, to concrete cellular behaviours essential for proper neural circuit formation.

## Discussion

In this study, we leveraged single-cell RNA sequencing of the E4 quail dorsal NT to construct a dI1-focused atlas and examine how transcriptional programs associated with this lineage change under RA inhibition. At this single yet critical developmental stage, our data capture a continuous trajectory from progenitor proliferation to the onset of neuronal migration, characterized by combinatorial sets of known ^5,7^ as well as novel genes whose functions await to be elucidated. Along with expression of unique markers, differentiated dI1 neurons also show a convergence toward a pan-neuronal gene profile shared with other dorsal interneuron subtypes ^7^. This developmental continuum is validated by RNA velocity and latent time analyses, and spatial gene expression by in situ hybridization confirms this developmental sequence along the medio-lateral extent of the NT ^8^.

Pathway enrichment and gene-expression patterns indicate that early dI1 states are dominated by cell-cycle and progenitor maintenance programs exemplified by *SOX2, SOX9* and *PAX7*, together with Notch pathway components. Genes associated with cytoskeletal organization, ECM remodeling and cell migration are induced in waves along the pseudotime, providing a molecular framework for the morphological changes that accompany neurogenesis. Apoptosis-related genes are transcriptionally enriched at early and late dp1, yet caspase-3 staining shows no significant apoptosis in the NT at E4–E4.5, implying a primed but inactive program, or non-apoptotic functions for these pathways ^76–78^. Their tightly staged expression peaks also argue against dissociation-induced stress as the cause of apoptotic gene upregulation.

Neuronal identity and function emerge through a temporally ordered sequence in which broad dorsal patterning genes give way to subtype-restricted determinants and, finally, to genes associated with mature neuronal signaling and connectivity. These patterns largely recapitulate principles established in vertebrate spinal cord development ^79^, and our dI1-focused analysis adds detail by mapping these transitions within a single dorsal interneuron class and comparing them to dI2 and dI4 trajectories.

Our data also reveal dynamic regulation of signaling pathways along the dI1 trajectory. BMP signaling, known to be crucial for dI1 patterning, is enriched in early progenitors, marked by expression of *ID1/3, MSX1/2* and pSmad activity, and progressively antagonized by inhibitors such as *HIPK2* ^80^, *FSTL4/5, CHRDL1 and DLX1* ^44,81^ as differentiation proceeds.

*RSPO2* and *RSPO3*, which can promote BMP receptor degradation ^69,82^, are enriched in differentiated dI1 neurons and are upregulated under RA inhibition, providing an additional layer by which BMP responsiveness might be curtailed. In contrast, at early progenitor stages, RA enhances expression of direct BMP genes, consistent with BMP stimulating cell proliferation as well as dI1 specification, together highlighting stage-specific functions of these morphogens in dI1 formation.

The above-mentioned antagonistic interactions of BMP signaling and its repressors is a recurring motif in dorsal NT development. Previous results highlighted the cross-talk between BMP and noggin in regulating NC delamination ^24,36,83^. Additionally, during the transition from NC to RP stages, BMP inhibitors such as *BAMBI*, *GREM1* and *RSPO3* are upregulated in the nascent RP, causing a loss of responsiveness to BMP and completion of NC emigration ^9,26^. We speculate that loss of BMP responsiveness in post-specification dI1 cells may also relate to changes in primary cilia, analogous to mechanisms described in ventral NT in which cilium loss renders delaminated cells insensitive to Shh, thereby promoting differentiation ^84,85^, but this remains to be tested experimentally.

The development of functional tissues and organs relies on extracellular signals ^86,87^. For example, RA exerts a positive effect on BMP signaling on the early progenitors (discussed above) and negatively modulates Wnt activity during neuronal differentiation, e.g; loss of the Wnt secreting protein *WLS* ^88^, of the Wnt agonist *RSPO1* ^89,90^, and/or by upregulation of the Wnt inhibitor *DACT1* ^53^, all observed in our data. In this context, it is relevant to mention that the BMP and RA-induced gene *BRINP1*, expressed along the dI1 trajectory, is repressed by RA primarily in proliferating progenitors and to a lesser extent in differentiating neurons. Since BRINP1 acts both as a cell cycle inhibitor and repressor of neurogenesis ^48,70^, we speculate that RA, via *BRINP1* repression, may enable the above processes to proceed at early and late stages, respectively.

Dorsal interneurons migrate long distances along the dorso-ventral axis to reach their distinctive locations and integrate into specific circuits. DI1 interneurons localize in the intermediate spinal cord and their proper positioning is crucial for their integration into neuronal circuitry ^8,34^. We find that loss of RA signaling compromises the normal positioning of dI1 interneurons. Already at E4, a subset of dI1 interneurons aberrantly infiltrates the RP domain, failing to segregate from NC and RP cells ^9,26^. At this stage, transcriptional changes in genes related to cell adhesion and cytoskeletal dynamics are already evident. Two days post-RA inhibition, BARHL2⁺ dI1 neurons showed significantly reduced ventral localization. By E6 (1 day later), when control interneurons had reached target zones, RA-deprived dI1 populations occupied ectopic territories ventral to their typical destinations. Additionally, at both stages, the number of dI1 neurons was significantly higher than in controls. This complicates the distinction between ectopic neuronal positioning due to impaired migration, increased neurogenesis, or a combination of both. Nevertheless, our data support the interpretation that dorsal accumulation primarily reflects compromised migration. First, the increase in the proportion of cells abnormally retained in dorsal locations surpasses what would be expected from the increase in total BarHL2^+^ dI1 neurons. Second, dI1 and dI3 neurons, whose ventral localization is similarly affected, exhibit opposite correlation patterns between total cell number and dorsal retention. Future experiments specifically designed to uncouple proliferation from migration would be required to dissect the primary cause of this later defect.

Despite these limitations, our single-timepoint dI1 atlas, combined with RA perturbation, provides a coherent framework for thinking about how transcriptional and signaling programs evolve within a dorsal interneuron lineage and how RA may intersect with these programs. We report that RA deprivation causes changes in genes and pathways regulating multiple developmental programs. For instance, upregulation of *BARHL1* and *BARHL2* in RARα403-treated embryos might be associated with neuronal differentiation and axon guidance ^16,66,91^ and is consistent with a general upregulation of terminal differentiation genes. RA also modulates Eph–Ephrin and Semaphorin–Plexin signaling ^92–97^, accompanied by stage-dependent downregulation of specific adhesion and cytoskeletal components. Although the precise genes involved in each function as well as the responsible mechanisms remain unresolved, these data provide a framework for dissecting RA-regulated development of dI1 interneurons.

Overall, our atlas reveals coordinated transcriptional and signaling programs underlying dI1 specification, differentiation, and migration from a single developmental stage, and highlights RA as both a temporal regulator and spatial organizer of spinal sensory circuit assembly.

## Experimental Procedures

### Experimental Model

Fertilized quail (Coturnix coturnix Japonica) eggs were obtained from commercial sources (Moshav Mata), kept at 15°C and then incubated at 38°C to desired stages. All experiments were performed on embryos younger than E6 and were therefore not subjected to IACUC regulations.

### Expression vectors and in-ovo electroporation

The following expression vectors were used: control pCAGG, pCAGGS-eGFP ^98^, and pCAGGS-RARα403 ^26^.

For NT electroporation, DNA (4 ug/ul) was mixed with Fast Green and microinjected into the lumen of the NT at the flank level of the axis. Five mm tungsten electrodes were placed on either side of the embryo. A square wave electroporator (BTX, San Diego, CA, USA) was used to deliver one pulse of current at 17 V for 6ms.

### NT collection, single-cell RNA sequencing and analysis

Single-cell RNA-seq data utilized in this study were previously published ^9^ and are accessible through GEO Series (See data availability). Briefly, embryos were electroporated at E2.5 with either control or RARα403 vectors, together with a Msx1-Citrine plasmid. At E4, ten NTs per group (with associated mesoderm) were mechanically microdissected and dissociated in 1.5 ml Accutase (Sigma Israel, A6964) containing 30 ul Papain (Sigma Israel, P3125) and 300 ng DNaseI (Sigma Israel, DN25). Samples were sorted using a BD FACSAria III sorter (BD Biosciences UK). Sorted cells were then subjected to droplet-based scRNA-seq (10x Genomics Chromium) to generate a gene expression profile of control and treated dorsal NT cells. In total, 17,549 cells were sequenced. After applying quality filters, a dataset of 14,566 cells was retained for further analysis (5,890 control and 8,676 treated cells, which comprise 86.6% and 80.7% of the initial number of cells, respectively).

Libraries were prepared following manufacturer’s instructions (GEM Single Cell 3ʹ GEM, Library & Gel Bead Kit v3.1, 10× Genomics, CA, USA). Approximately 10.000 cells were loaded per sample. Sequencing was performed using the Illumina Nextseq500 platform (Illumina Inc, San Diego, CA, USA). As detailed in the original publication, reads were aligned and quantified using the Cell Ranger pipeline ^99^ (v6.0.1, 10x Genomics) against the *Coturnix coturnix Japonica* 2.1 genome (RefSeq assembly GCF_001577835.2). The gene annotation was filtered using Cell Ranger’s mkgtf command to keep protein-coding genes and lncRNAs. Downstream analysis and visualization were performed using the Seurat R package ^100^ (v4.04).

To identify cluster-specific marker genes, differential expression analysis was performed using Seurat’s ‘FindAllMarkers’ function, which applies the non-parametric Wilcoxon rank-sum test to compare gene expression in each cluster against all other clusters. This approach systematically pinpoints genes significantly upregulated in a given cluster relative to the remaining cell groups. Only the control sample was used for marker identification. Cell types were assigned manually based on the expression of characterized genes^9^

Seurat’s functions ‘FeaturePlot’ and ‘DimPlot’ were used for visualization. Seurat’s ‘DotPlot’ function was used to generate dot plots. Plots were further formatted using custom R scripts with ggplot2. Heatmaps were produced with The R package ComplexHeatmap (Version 2.14)^103^

Out of the 16,706 genes in our data, 4422 had uncertain function (LOC symbols; LOC plus the GeneID). Of them, 76 LOC genes that exhibited notable expression patterns, were assigned names based on high similarity to orthologous genes (Available through Mendeley Data Repository, see Data Availability).

### RNA Velocity, Latent time and Trajectory analysis

Velocyto (version 0.17.17) ^104^ was used to generate count matrices, based on the spliced and unspliced reads of the RARα403-treated cells, using the ‘run10x’ option. Unspliced and spliced counts were matched to the barcodes and genes retained after filtering. scVelo (version 0.2.5), an unbiased approach that leverages the distinction of unspliced and spliced RNA transcripts from the aligned sequences ^100^, was used to compute velocities using the following options: ‘filter_and_normalize’ was run using ‘min_shared_counts’ set to 20 and ‘min_cells’ set to 80; ‘monemts’ was run with ‘n_neighbors’ set to 10. Velocities were calculated using the ‘dynamical’ mode and visualized on the UMAP. Latent time was calculated using scVelo ‘tl.latent_time’ function. The RNA velocity analysis was performed without regressing out the cell cycle effect. Trajectory inference and pseudotime estimation were performed using Slingshot (v2.7.0) ^105^.

Gene filtration for Figure 2B – Genes changing significantly along the control dI1 trajectory were identified using two complementary approaches: **1) Cluster-specific marker identification**: For each of the four dI1 clusters, we performed differential expression analysis comparing it against the combined cells from the other three dI1 clusters using the non-parametric Wilcoxon rank sum test (Seurat’s ‘FindAllMarkers’ function). Genes were included if they had a linear fold change above 1.5, adjusted p-value below 0.05, and were expressed in at least 50% of the cells in the cluster. **2) Pairwise cluster comparison**: We performed differential expression analysis between all possible pairs of the four dI1 clusters. Genes with a linear fold change above 1.5 and adjusted p-value below 0.05 were considered differentially expressed.

To enrich for specific expression patterns, we excluded 207 ubiquitously expressed genes (those present in more than 60% of both the entire control sample and dI1 cells, unless their expression in the control sample was below 70%). A total of 954 genes passed these combined criteria and are included in Figure 2B and the Mendeley Data Repository (See Data Availability).

Gene filtration for Sup. Figure 14 (differentially expressed genes) is explained in the corresponding legend. Data is available through the Mendeley Data Repository (see Data Availability).

### Functional pathways - Specific gene curation

Gene sets aligned with biological pathways were systematically compiled for the control arm of the dI1 transcriptome. Pathway enrichment analysis was performed using Enrichr (using the Reactome database) ^106^ to detect pathways (version 2024, doi: 10.1093/nar/gkad1025) significantly associated with genes exhibiting significant expression changes in each cluster along the dI1 developmental trajectory (Figure 2B). Manual curation was then applied to include additional relevant genes whose functions, as described in the literature, align with these pathways. Some pathways were combined to avoid redundancy.

To identify functional pathways in differentially expressed genes between control and RARα403-treated embryos, genes were manually curated based on published data. Pathway average score per cell was calculated by averaging the normalized data of the pathway genes.

The functional pathway lists are available through Mendeley Data Repository, (see Data Availability).

Scatter plots with the Pathway Average score lines and cell cycle data - In the scatter plots, each dot represents a cell. The pathway score is plotted as a function of the pseudotime. Cells are colored according to their predicted cell cycle phase (calculated by Seurat’s CellCycleScoring function). A LOESS-smoothed trend line was fitted to the data using the geom_smooth function from ggplot2.

### Immunohistochemistry

For immunostaining, embryos were fixed overnight at 4°C with 4% formaldehyde (PFA) in PBS (pH 7.4), embedded in paraffin wax and serially sectioned at 8 μm. Immunostaining was performed on whole mounts or paraffin sections, as previously described ^101,102^. Antibodies were diluted in PBS containing 5% fetal bovine serum (Biological Industries Israel, 04-007-1A) and 1% or 0.1% Triton X-100 (Sigma Israel, X-100), respectively. Antibodies used were: rabbit anti-pSmad1/5/9 (1:500, Cell Signaling Technology, CST13820), rabbit anti-Cleaved Caspase3 (1:200, Cell Signaling Technology, CST9661), rabbit anti-GFP (1:1000, Invitrogen, Thermo-Fisher Scientific, A-6455), chicken anti-GFP (1:500, Novus, NB100-1614), rabbit anti-BarHL1 (1:300, Sigma Israel, HPA004809), rabbit anti-BarHL2 (1:100, Sigma Israel 23976-a-AP), mouse anti-Brn3a (1:300, Millipore MAB1585), rabbit anti-Sox2 (1:300, Abcam, ab92494), mouse anti-Isl1 (1:50, DSHB, 4D5) and mouse anti-Pax7 (1:10, DSHB, PAX7), mouse anti-H3-pS10 (1:400, Abcam, Ab14955). Cell nuclei were visualized with 125 ng/ml Hoechst 33258 (Sigma Israel, 14530) diluted in PBS.

### In situ hybridization

For ISH, embryos were fixed in Fornoy (60% ethanol, 30% formaldehyde and 10% acetic acid) for 1 hr at room temperature, embedded in paraffin wax and serially sectioned at 10 μm. Briefly, sections were treated with 1 µg/ml proteinase K, re-fixed in 4% PFA, then hybridized overnight at 65 °C with digoxigenin-labeled RNA probes (Roche, 11277073910). The probes were detected with AP coupled with anti-digoxigenin Fab fragments (Roche, 11093274910). AP reaction was developed with 4-Nitro blue tetrazolium chloride (NBT, Roche, 11383213001) and 5-bromo-4-chloro-3’-indolyphosphate p-toluidine salt (BCIP, Sigma Israel, B8503). Importantly, all hybridizations were always developed for the same time for a specific probe and experiment. Probes were synthesized from PCR products (see Data Availability) using the KAPA2G Fast ReadyMix PCR kit, (Sigma Israel, KK5101). cDNA templates were synthesized by RNA precipitation followed by reverse transcription PCR. RNAs were produced from 20 somite-stage-to-E4 quail embryos. Tissue samples were homogenized with TriFast reagent, and RNA was separated with chloroform and isopropanol.

### Data acquisition, quantification and statistical analysis

Images were photographed using a DP73 (Olympus) cooled CCD digital camera mounted on a BX51 microscope (Olympus) with Uplan FL-N 20 x/0.5 and 40 x/0.75 dry objectives (Olympus).

For quantification, images of control and treated sections were photographed under the same conditions. Fluorescence/ISH signals were quantified in transverse sections. The number of embryos and sections monitored per treatment is depicted in the respective figure legends. Intensities of immunofluorescent and ISH signals were measured using FIJI software ^107^. The dorsal neural tube was delineated as the region of interest (ROI) for most analyses. For intensity measurements, mean signal intensity was calculated with background subtraction. For area measurements, specific staining was defined by applying consistent thresholds across all samples, with areas above threshold quantified. For each embryo, measurements were averaged across all sections (numbers of embryos and sections are indicated in respective figure legends). All intensity values were normalized to the control group mean, which was set to 1.

To analyze ventral migration, the distance of BarHL2^+^ cells from the dorsal midline of the NT was measured for each cell manually using FIJI. Statistical analysis and data visualization were performed using R in RStudio. Briefly, the length and angle of the distance measured, was converted to x and y coordinates and normalized to NT dimensions. To visualize spatial cell distributions, we applied 2D kernel density estimation (KDE) using the kde2d function from the R MASS package on a 100X100 grid. The KDE output was normalized to represent the percentage of total cells per grid cell. To define density contours, we sorted all grid values and calculated thresholds that enclosed specific cumulative percentages of the cell populations. Complete analysis scripts and raw data are available in Table S1 and the Mendeley Data repository (See Data Availability).

For figure preparation, images were exported into Photoshop CS6 (Adobe). If necessary, the levels of brightness and contrast were adjusted to the entire image. Graphics were generated using Graphpad Prism 9.0 and R version 4.4.2 (https://www.R-project.org/) in Rstudio (http://www.rstudio.com/). Scatter plots show mean ± SEM. Data were subjected to statistical analysis using either of the following tests, as described in the respective figure legends: Student’s t-test, Welch’s t-test, Mann-Whitney test and Kolmogorov-Smirnov test. All tests applied were two-tailed and a p-value ≤ 0.05 was considered significant. Figures were prepared using InDesign CS6.

## Acknowledgements

We express our gratitude to Avihu Klar for careful review of the manuscript. GF is the Incumbent of the David and Stacey Cynamon Research fellow Chair in Genetics and Personalized Medicine.

## Author Contributions

DR performed most experiments, analyzed the data, and wrote the manuscript. GF led the bioinformatic analysis assisted by DR and CK. SK prepared probes and assisted with ISH. NRK analyzed the data and performed ISH. SV contributed to cell cycle and migration experiments. CK conceived and supervised the project, analyzed the data and wrote the manuscript. All authors discussed and agreed on the text and approved the manuscript.

## Notes

**<u>Grant Sponsor</u>**: This study was supported by grants from the Israel Science Foundation (ISF #219/23 to CK). The funders had no role in study design, data collection and analysis, decision to publish, or preparation of the manuscript.

### Competing Interest Statement

The authors have declared no competing interest.

### Summary of Updates

The following revisions were made: 1)Functional data on cell proliferation performed.(new figure 5) 2) Further validations of gene expression performed by in situ hybridization (new figure 7) 3) Dotplot depicting the identification of dI1 interneurons (Fig.1) The order of figures was slightly changed as some were moved to the supplem section and vice-versa

