## Supplemental figures 1-16 for "Dynamic Functional Pathway Development in Type 1 Spinal Interneurons: Stage-specific roles of retinoic acid activity"

### Supplementary Figure Titles and Legends

**Supplementary Figure 1 – Molecular markers characteristic of the control dI1 cluster 2**

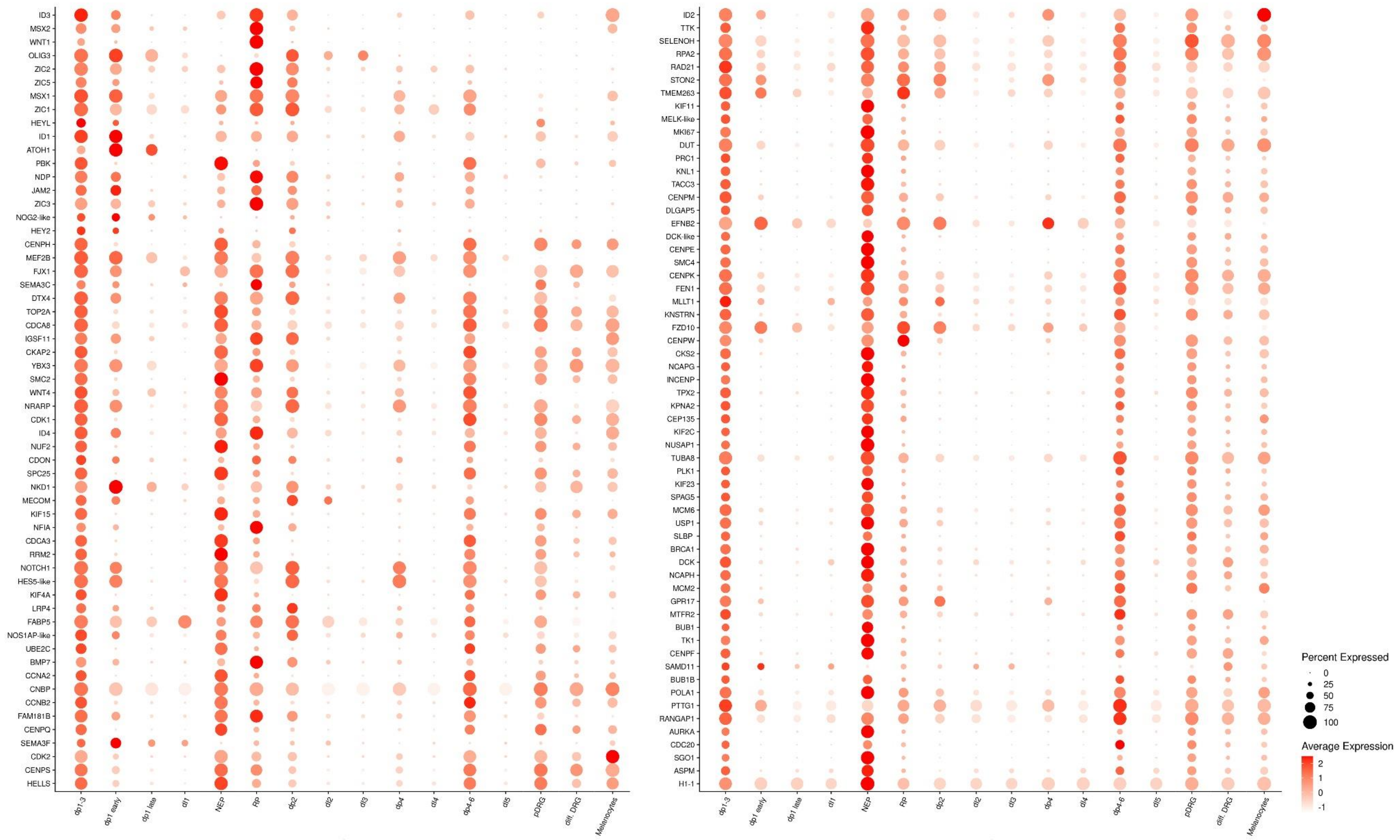

Dot plots visualizing selected marker gene expression in dI1 cluster 2 (dp1-3) relative to the remaining UMAP clusters. The size of the dot corresponds to the percentage of cells expressing the gene in each cluster. The color represents the average expression level. Genes were initially identified by differential expression analysis (linear fold change > 1.4, adjusted p-value < 0.05). A subset was curated manually using the 10X Loupe Browser to exclude genes with widespread or non-specific expression patterns.

**Supplementary Figure 2 – Molecular markers characteristic of the control dI1 cluster 8**

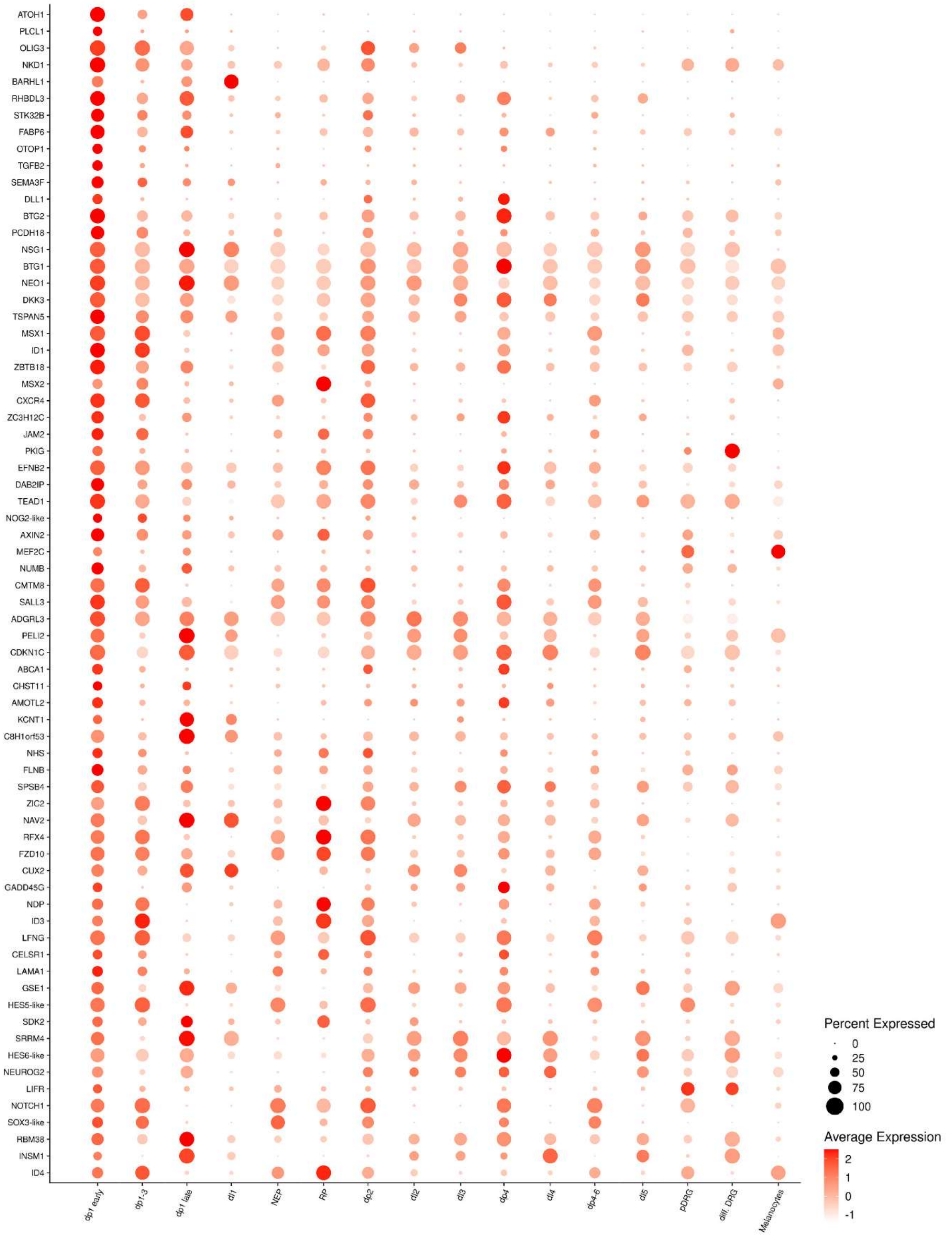


Dot plots visualizing selected marker gene expression in dI1 cluster 8 (dp1 early) relative to the remaining UMAP clusters. The size of the dot corresponds to the percentage of cells expressing the gene in each cluster. The color represents the average expression level. Genes were initially identified by differential expression analysis (linear fold change $> 1.4$, adjusted p-value $< 0.05$). A subset was curated manually using the 10X Loupe Browser to exclude genes with widespread or non-specific expression patterns.

#### Supplementary Figure 3 – Molecular markers characteristic of the control dI1 cluster 4

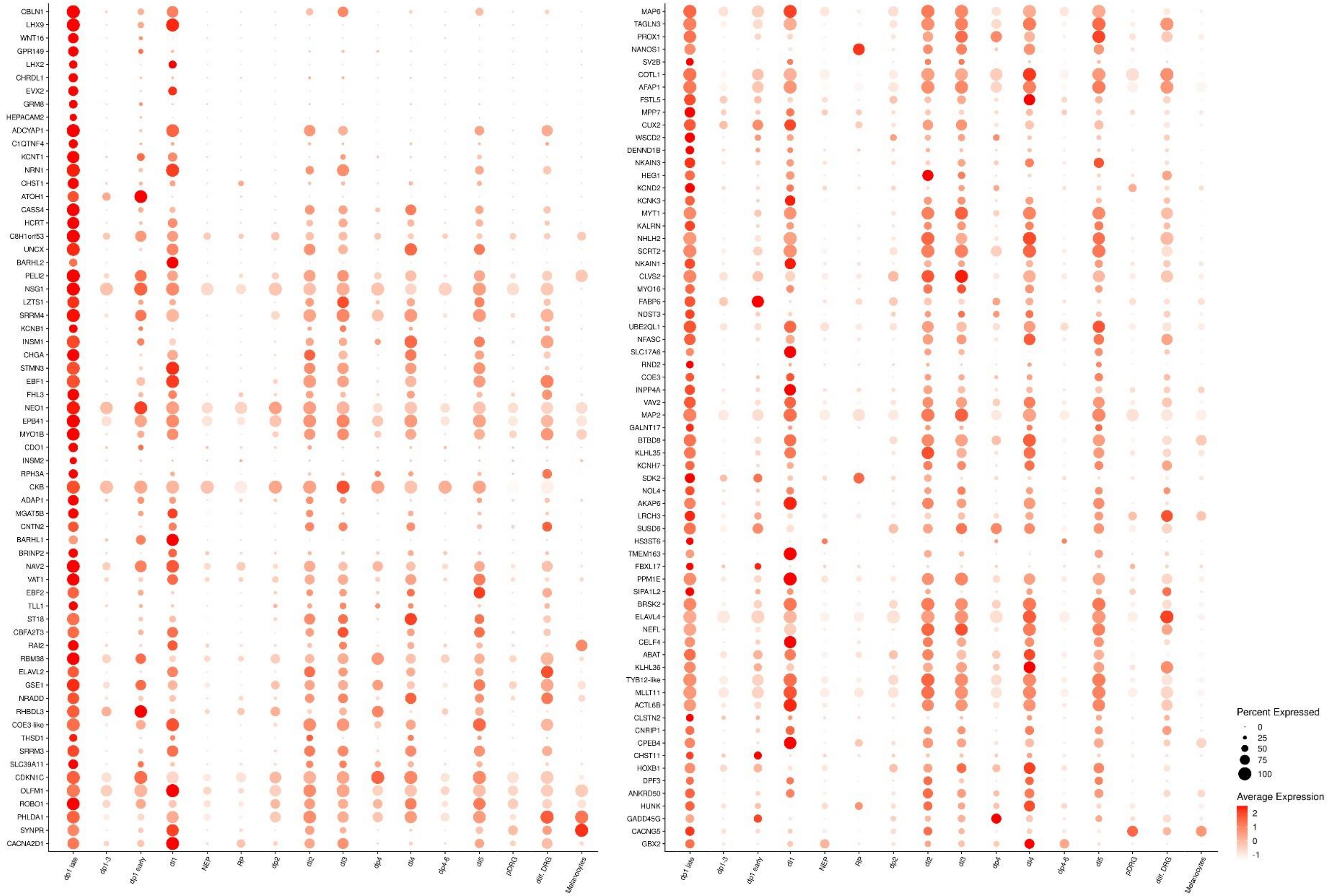

Dot plots visualizing selected marker gene expression in dI1 cluster 4 (dp1 late) relative to the remaining UMAP clusters. The size of the dot corresponds to the percentage of cells expressing the gene in each cluster. The color represents the average expression level. Genes were initially identified by differential expression analysis (linear fold change > 1.4, adjusted p-value < 0.05). A subset was curated manually using the 10X Loupe Browser to exclude genes with widespread or non-specific expression patterns.

#### Supplementary Figure 4 – Molecular markers characteristic of the control dI1 cluster 9

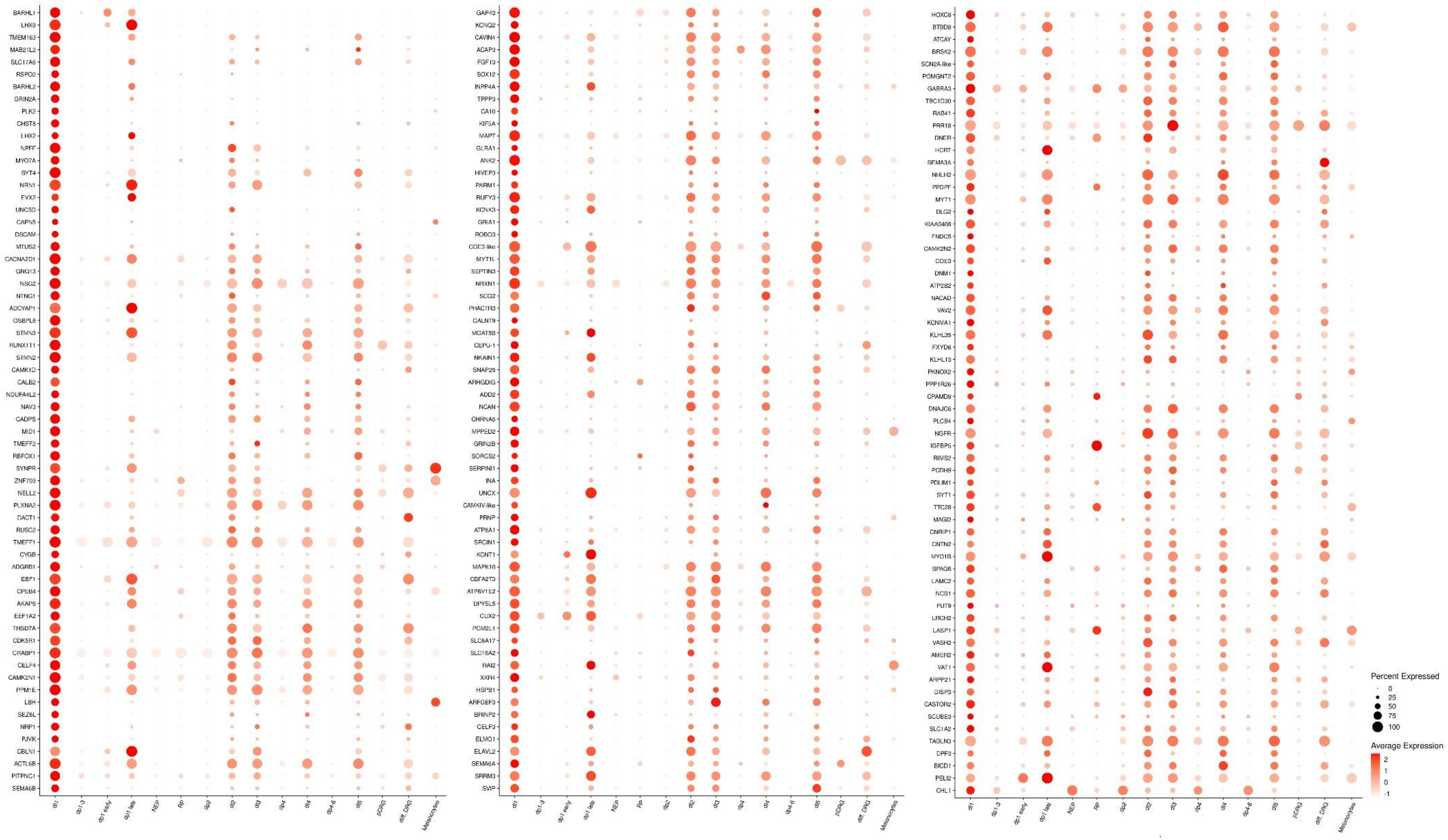

Dot plots visualizing selected marker gene expression in dI1 cluster 9 (dI1) relative to the remaining UMAP clusters. The size of the dot corresponds to the percentage of cells expressing the gene in each cluster. The color represents the average expression level. Genes were initially identified by differential expression analysis (linear fold change > 1.4, adjusted p-value < 0.05). A subset was curated manually using the 10X Loupe Browser to exclude genes with widespread or non-specific expression patterns.

**Supplementary Figure 5 - Dynamic regulation of genes associated with Mitosis and DNA Replication/Repair along the dI1 differentiation trajectory**

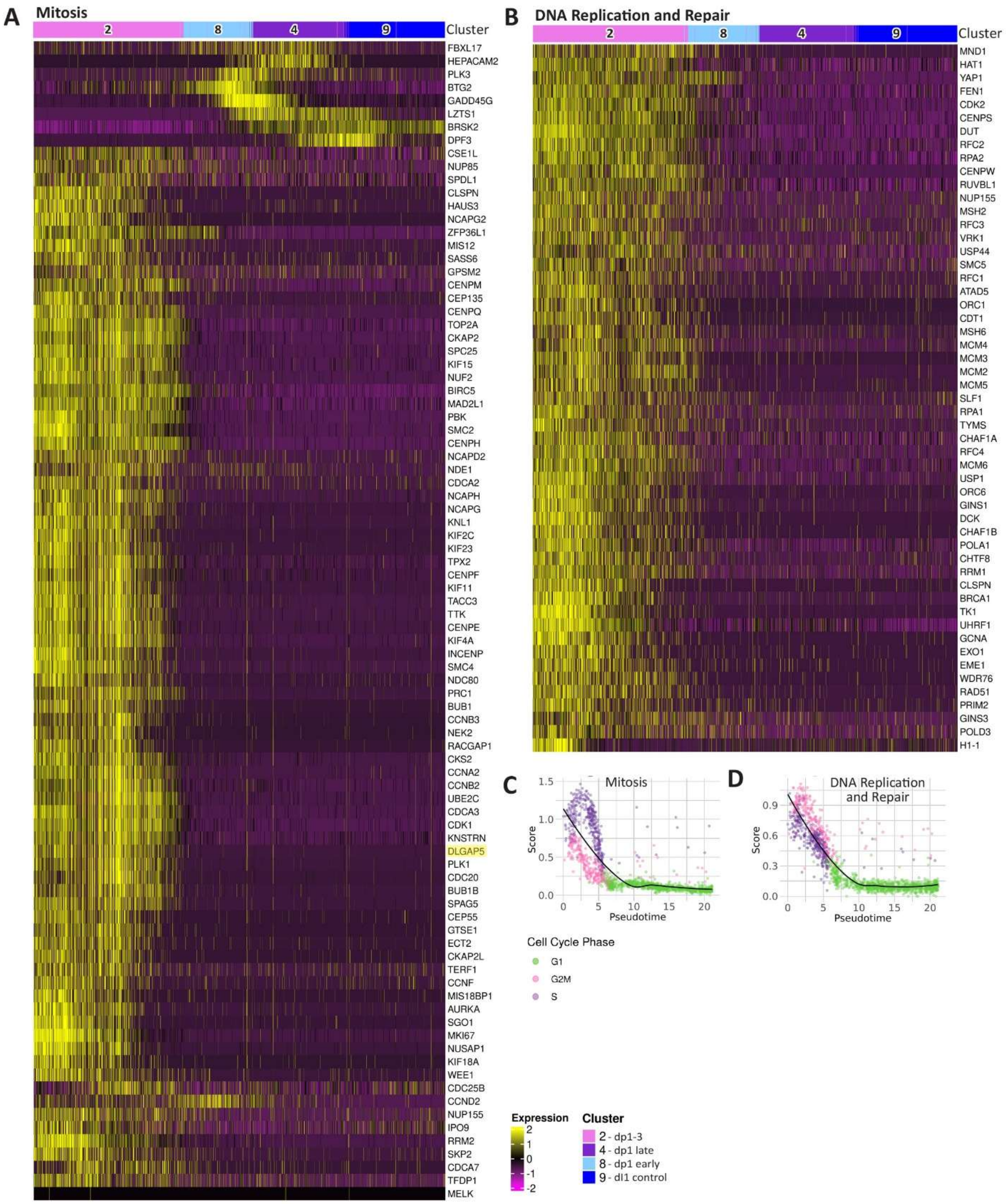


A-B) Heatmaps showing expression of genes associated with Mitosis and DNA Replication and Repair pathways along the dI1 control differentiation trajectory. DLGAP5 (validated in Figure 2D) is highlighted in yellow. Shown are Seurat scaled expression values. Cells are ordered by pseudotime, and the cluster number is indicated as top annotation.

C-D) Average gene expression levels of genes associated with the above pathways, along the pseudotime, with cells color-coded by cell cycle phase (See Methods).

**Supplementary Figure 6 - Temporal expression of genes associated with transcriptional regulation and inhibition of differentiation during dI1 development**

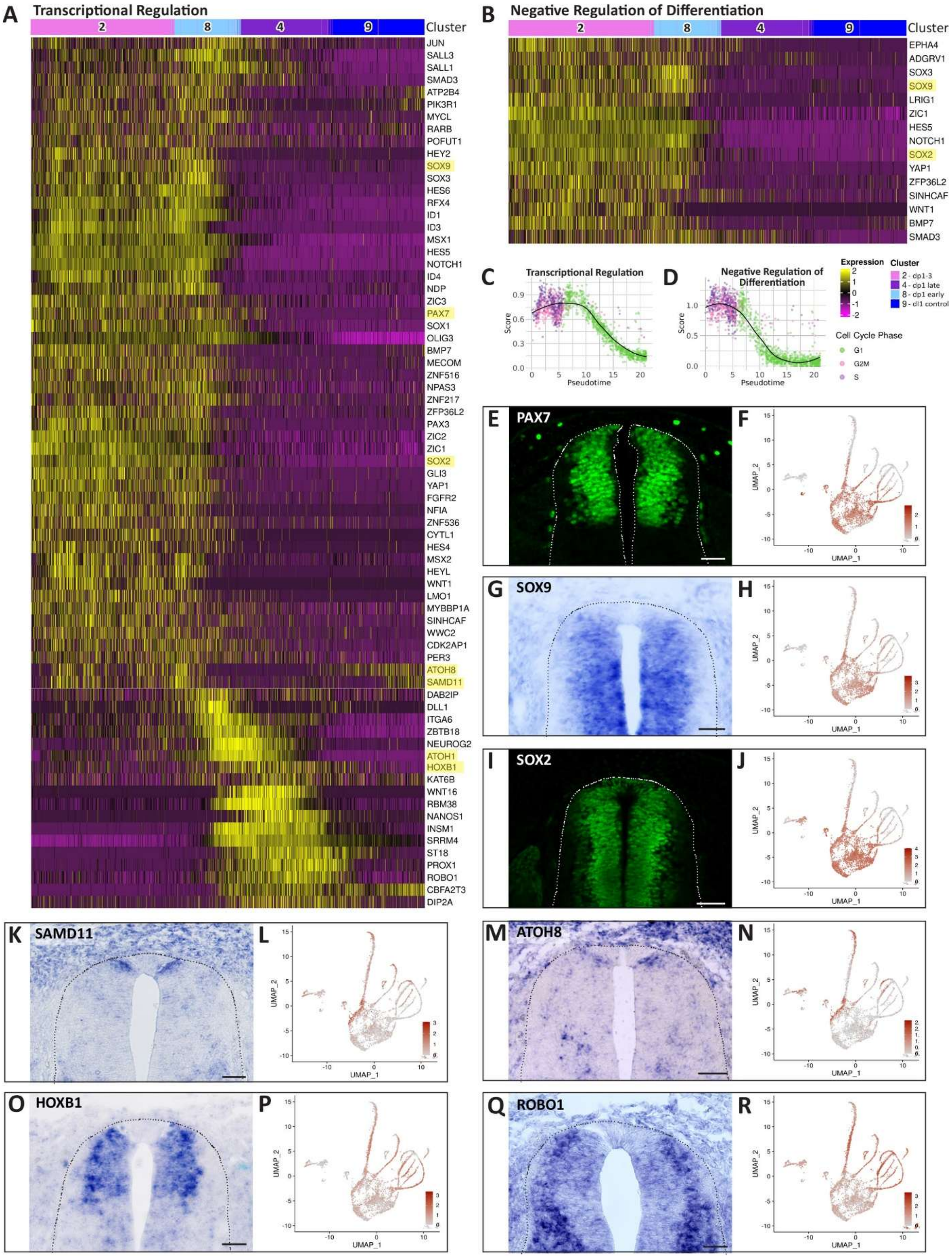

A-B) Heatmaps showing expression of genes associated with Transcriptional Regulation and Negative Regulation of Differentiation pathways along the dI1 control trajectory. Shown are Seurat scaled expression values. Validated genes (E-R) are highlighted in yellow. Cells are ordered by pseudotime, and the cluster number is indicated as top annotation.

C-D) Average gene expression levels of genes associated with the above pathways, along the pseudotime, with cells color-coded by cell cycle phase (See Methods).

E-R) ISH (G,K,M,O,Q) or immunostaining (E,I) of E4 embryos for representative genes of the above pathways, with corresponding UMAP projections of expression patterns F,H,J,L,N,P, R). Scale bar, 50 μm.

**Supplementary Figure 7 - Dynamic expression of Wnt, Notch, and Semaphorin-Plexin signaling pathway genes along the dI1 differentiation trajectory**

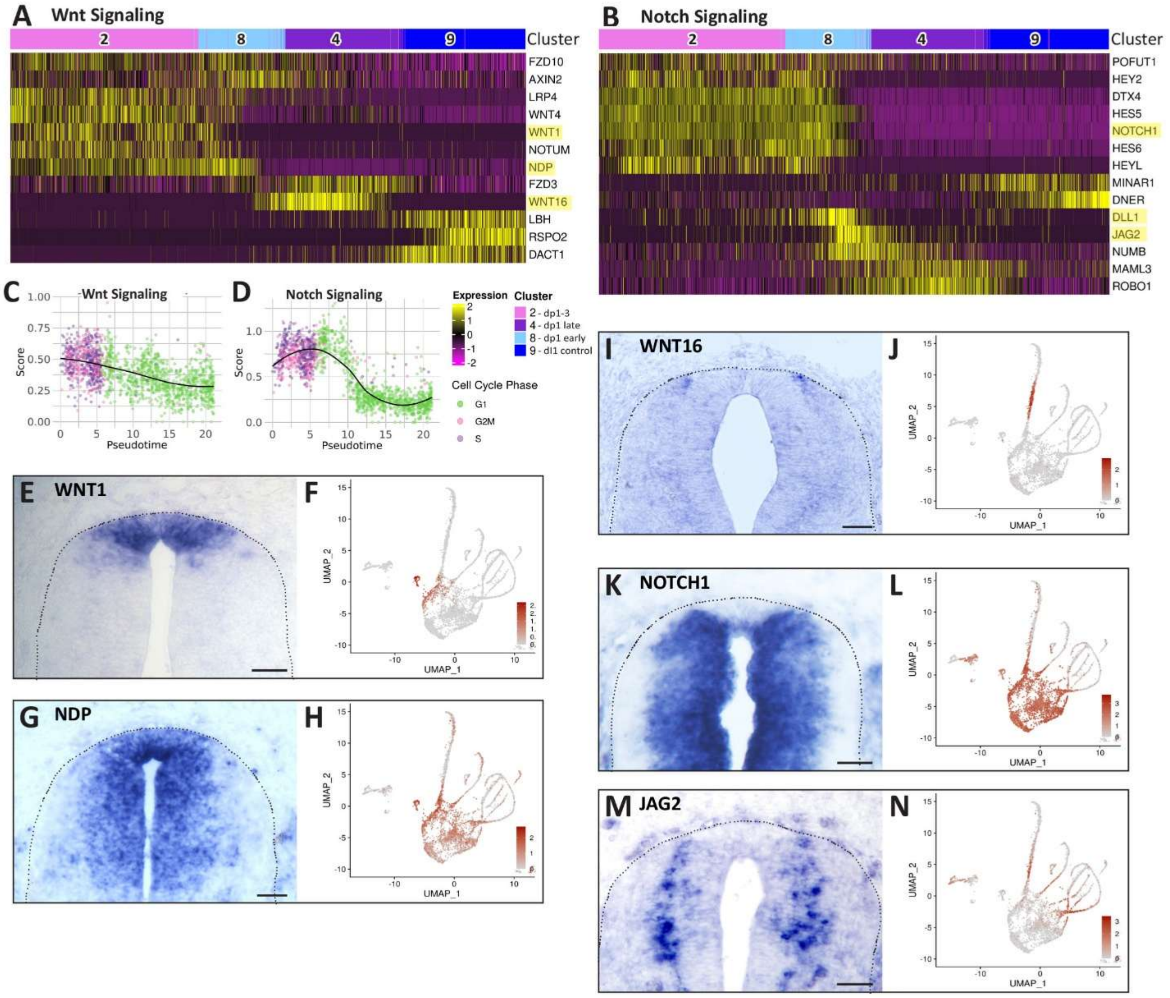


A-B) Heatmaps showing expression of genes associated with the Wnt and Notch signaling pathways along the dI1 control trajectory. Shown are Seurat scaled expression values. Validated genes (E-N and Figure 2B) are highlighted in yellow. Cells are ordered by pseudotime, and the cluster number is indicated as top annotation.

C-D) Average gene expression levels of genes associated with the above pathways, along the pseudotime, with cells color-coded by cell cycle phase (See Methods).

E-N) ISH for representative genes of the above pathways in E4 embryos, with corresponding UMAP projections of expression patterns.

Scale bar, 50 μm.

**Supplementary Figure 8 - Coordinated regulation of cytoskeletal dynamics, ECM remodeling, and migratory programs during dI1 interneuron development**

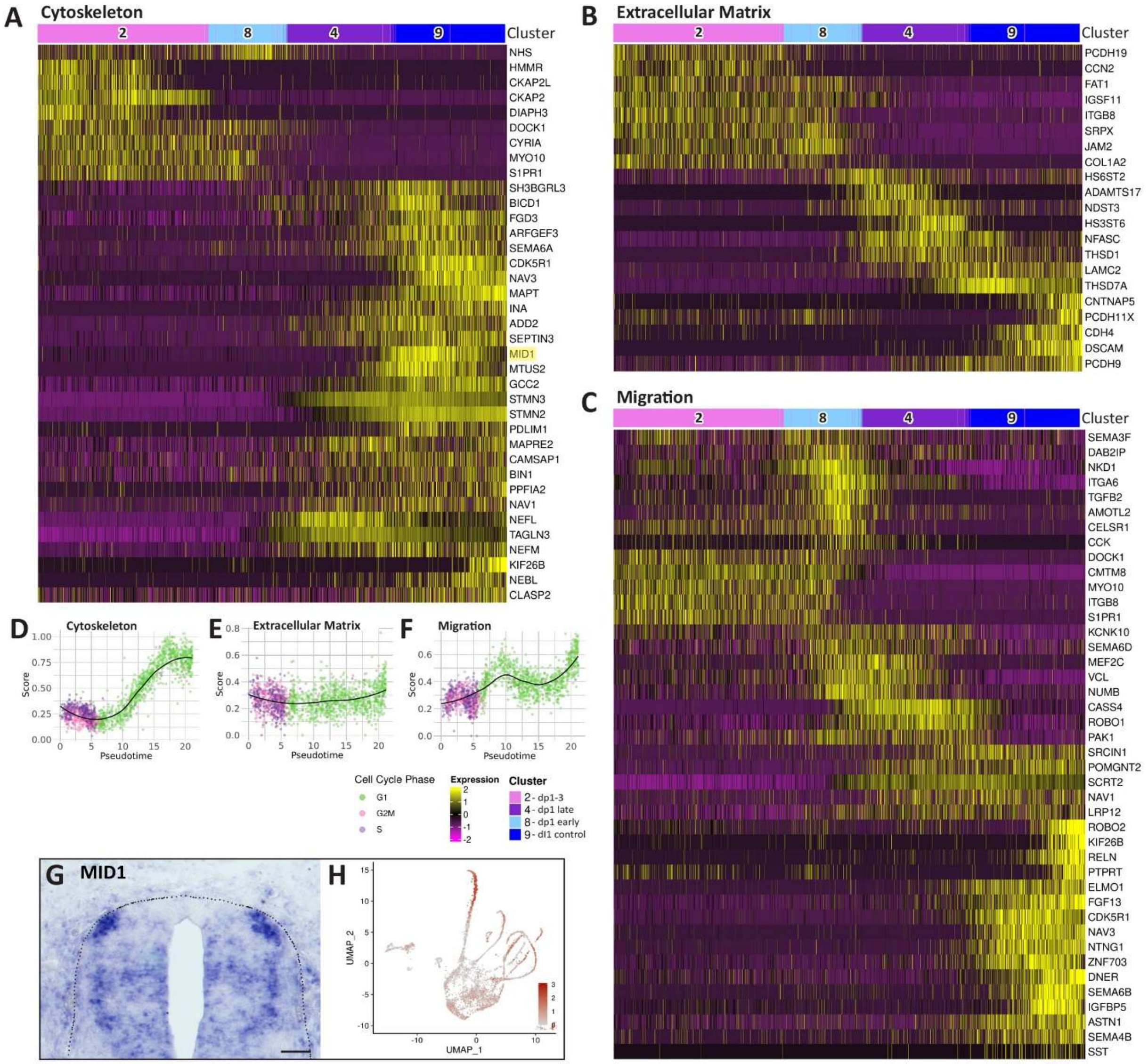


A-C) Heatmaps showing expression of genes associated with the Cytoskeleton, Extracellular Matrix and Migration pathways along the dI1 control trajectory. Shown are Seurat scaled expression values. *MID1* (validated in G) is highlighted in yellow. Cells are ordered by pseudotime, and the cluster number is indicated as top annotation.

D-F) Average gene expression levels of genes associated with the above pathways, along the pseudotime, with cells color-coded by cell cycle phase (See Methods).

G-H) ISH for *MID1* of an E4 embryo, with a corresponding UMAP projection of the expression pattern.

Scale bar, 50 μm.

**Supplementary Figure 9 – Neuronal Specification, Differentiation and Signaling are sequentially activated during dI1 interneuron development**

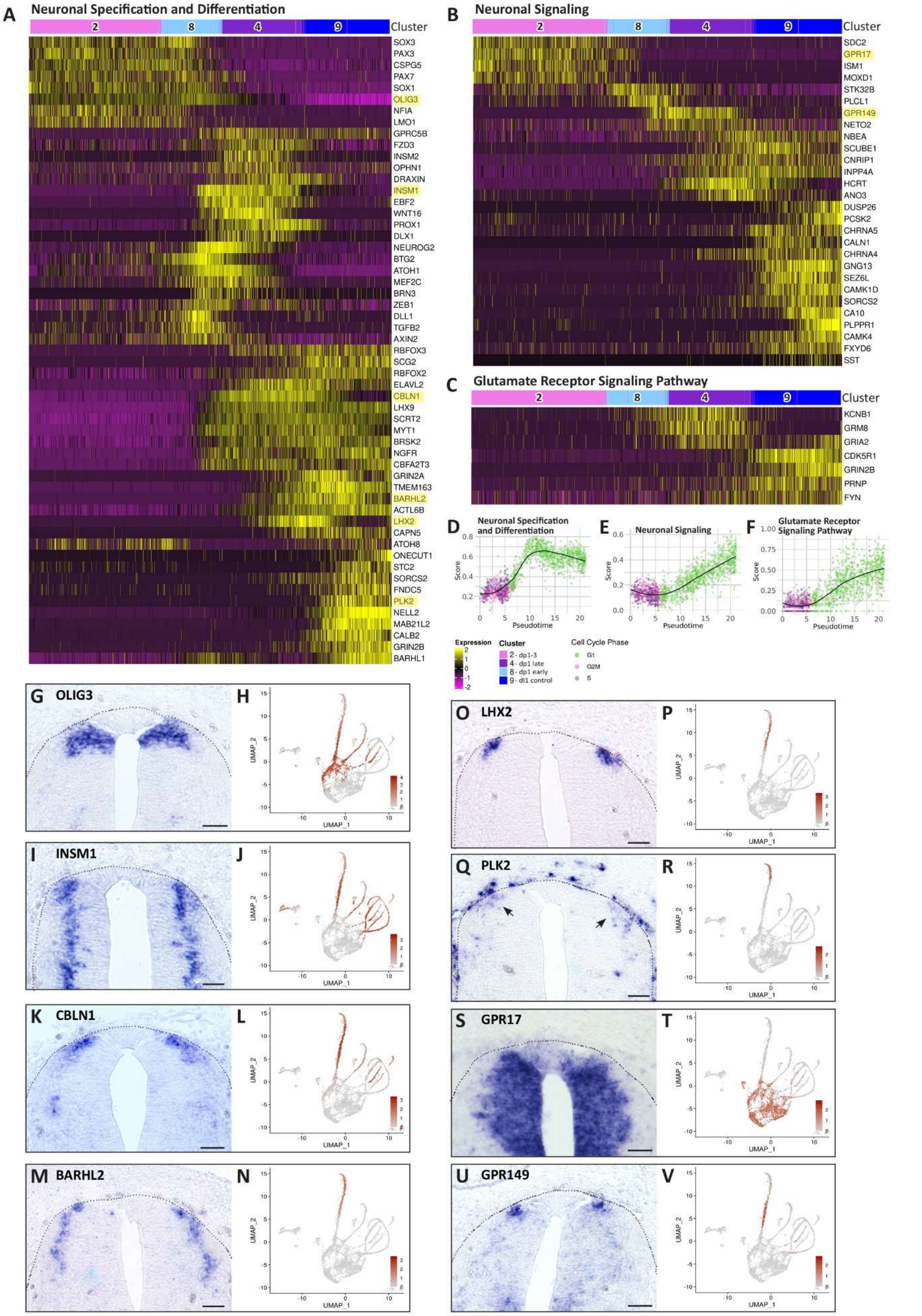

A-C) Heatmaps showing expression of genes associated with Neuronal Specification and Differentiation, Neuronal Signaling and Glutamate Receptor Signaling, along the dI1 control trajectory. Shown are Seurat scaled expression values. Validated genes (G-V) are highlighted in yellow. Cells are ordered by pseudotime, and the cluster number is indicated as top annotation.

D-F) Average gene expression levels of genes associated with the above pathways, along the pseudotime, with cells color-coded by cell cycle phase (See Methods).

G-V) ISH for representative genes of the above pathways in E4 embryos, with corresponding UMAP projections of expression patterns.

Scale bar, 50 μm.

**Supplementary Figure 10 – Neuronal Projections and Ion Transport Programs are activated in the final stages of dI1 interneuron differentiation**

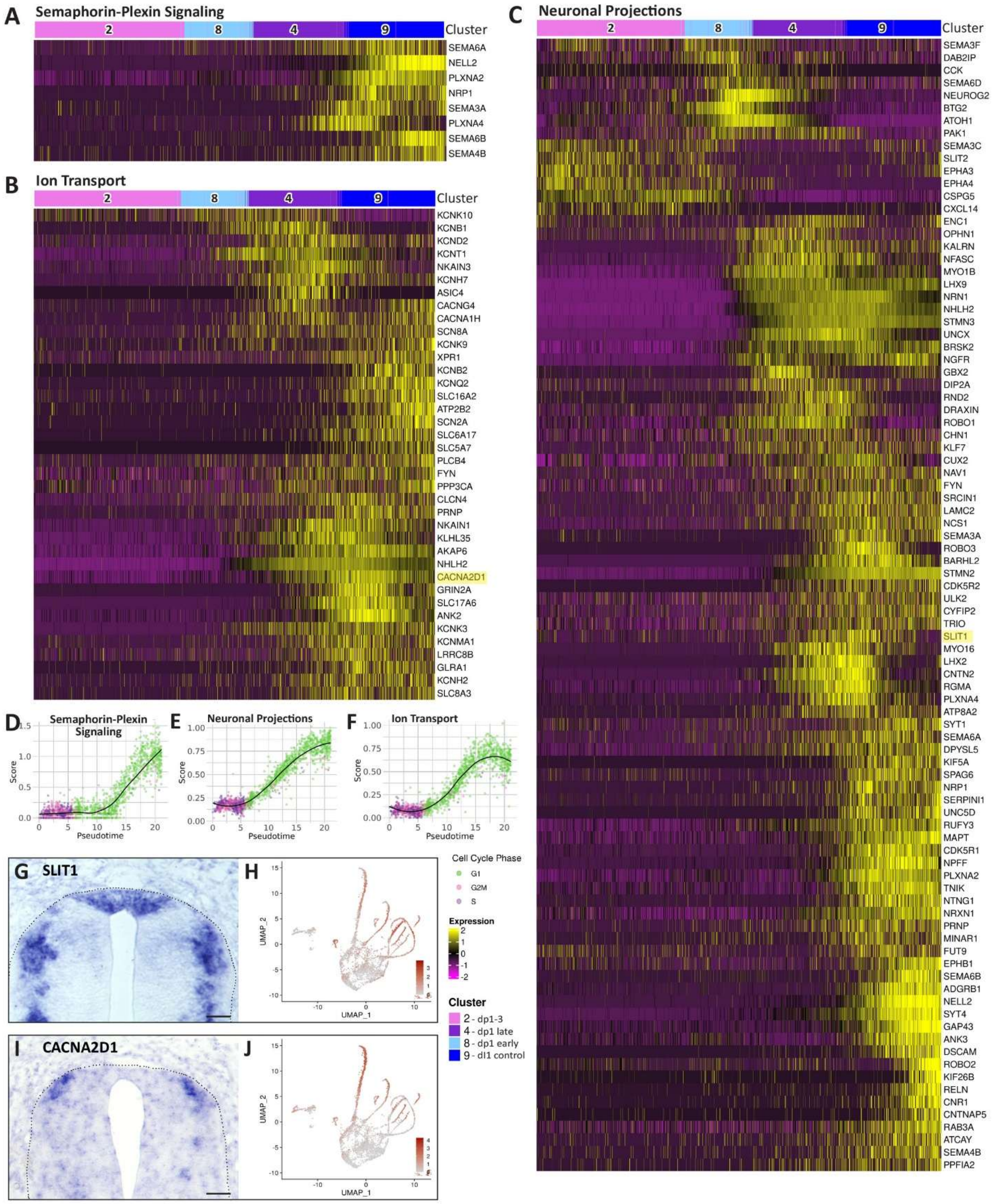


A-C) Heatmaps showing expression of genes associated with Semaphorin-Plexin signaling, Neuronal Projections and Ion Transport pathways along the dI1 control trajectory. Validated genes (G-J) are highlighted in yellow. Cells are ordered by pseudotime, and the cluster number is indicated as top annotation.

D-F) Average gene expression levels of genes associated with the above pathways, along the pseudotime, with cells color-coded by cell cycle phase (See Methods).

G-J) ISH for representative genes of the above pathways in E4 embryos, with corresponding UMAP projections of expression patterns.

Scale bar, 50 μm.

**Supplementary Figure 11 - Synaptic Transmission, Synapse Assembly, and Vesicle Trafficking Pathways are induced during an advanced stage of dI1 interneuron development**

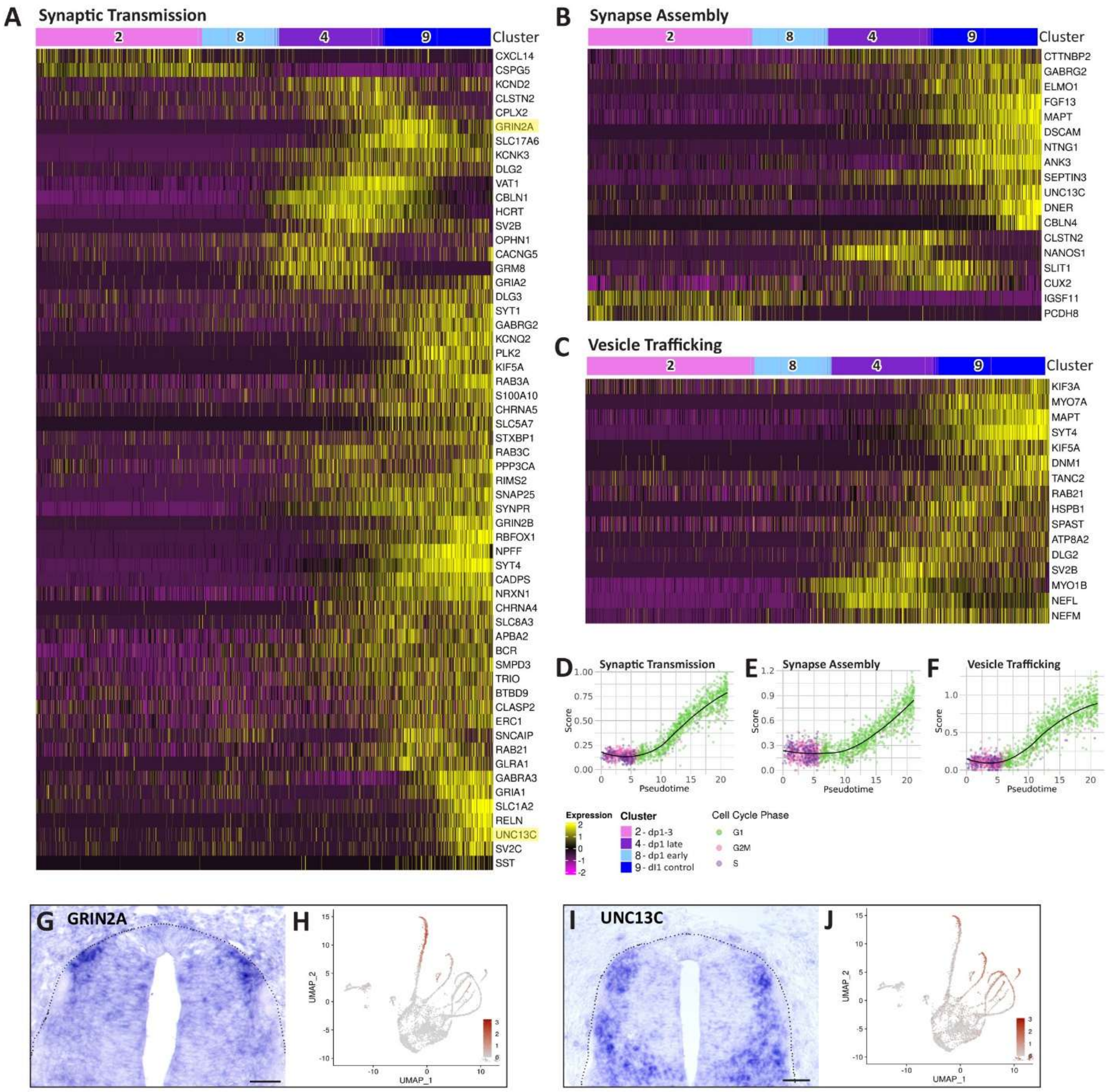


A-C) Heatmaps showing expression of genes associated with the pathways Synaptic Transmission, Synapse Assembly and Vesicle Trafficking, along the dI1 control trajectory. Validated genes (G-J) are highlighted in yellow. Cells are ordered by pseudotime, and the cluster number is indicated as top annotation.

D-F) Average gene expression levels of genes associated with the above pathways, along the pseudotime, with cells color-coded by cell cycle phase (See Methods).

G-J) ISH for representative genes of the above pathways in E4 embryos, with corresponding UMAP projections of expression patterns.

Scale bar, 50 μm

**Supplementary Figure 12 – Comparison of transcriptional dynamics between dorsal interneuronal populations**

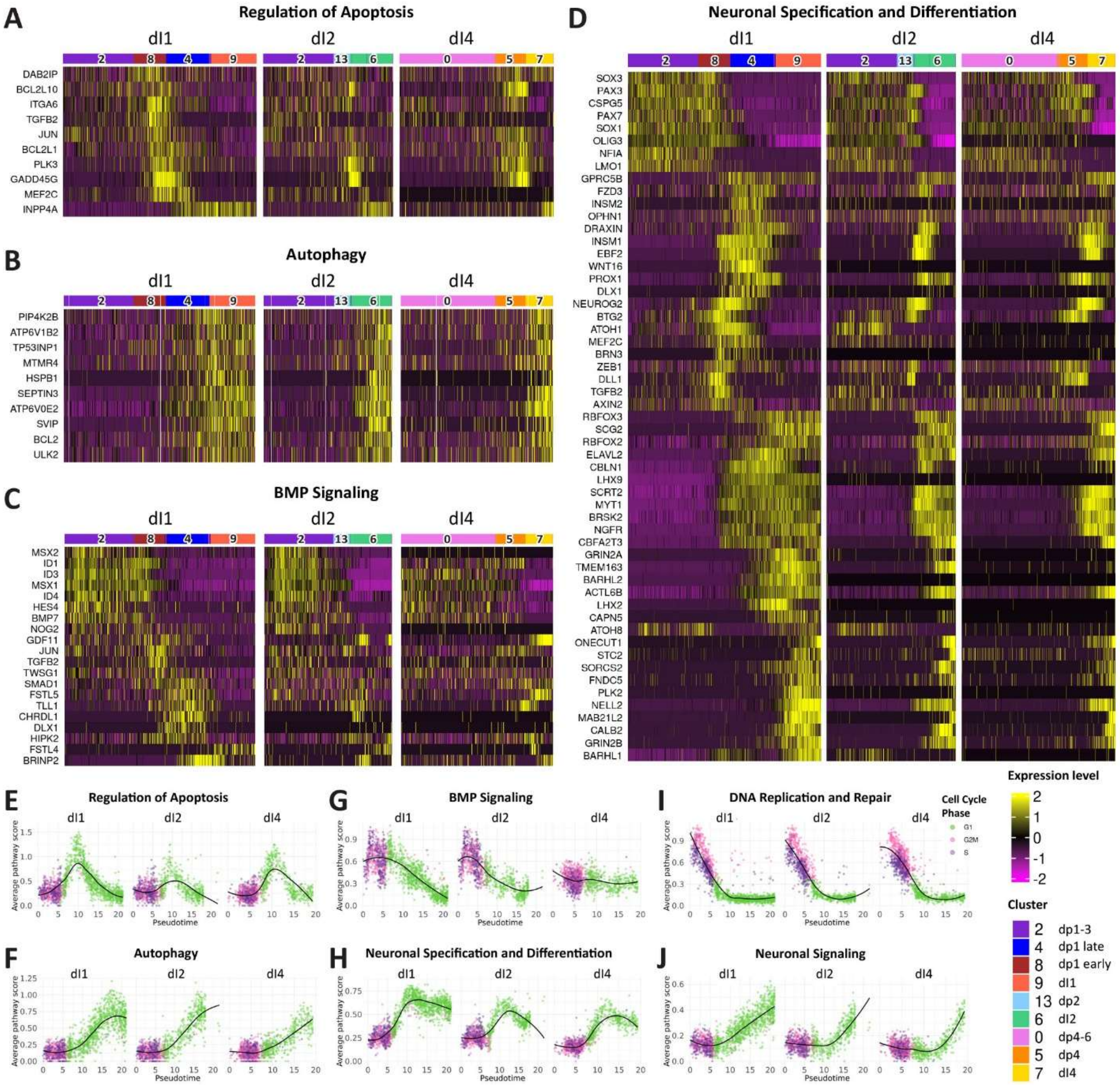


Trajectory inference and pseudotime estimation were performed for the dI1, dI2 and dI4 control cell populations using Slingshot (See Methods). Several functional pathways analysed in dI1 cells were additionally tested in dI2 and dI4 populations, using the same gene lists.

A-D) Heatmaps of genes associated with selected pathways in control cells: (A) Apoptosis; (B) Autophagy; (C) BMP Signaling and (D) Neuronal Specification and Differentiation. Shown are Seurat scaled expression values. Cells are ordered by the pseudotime, and the cluster number is shown as top annotation.

(E-J) Average expression levels of genes of selected pathways, as a function of the pseudotime, with cells color-coded by cell cycle phase: (E) Regulation of Apoptosis; (F) Autophagy; (G) BMP Signaling; (H) Neuronal Specification and Differentiation; (I) DNA Replication and Repair; (J) Neuronal Signaling.

**Supplementary Fig. 13. Control and RARα403-treated cells exhibit shared dI1 interneuron differentiation trajectories**

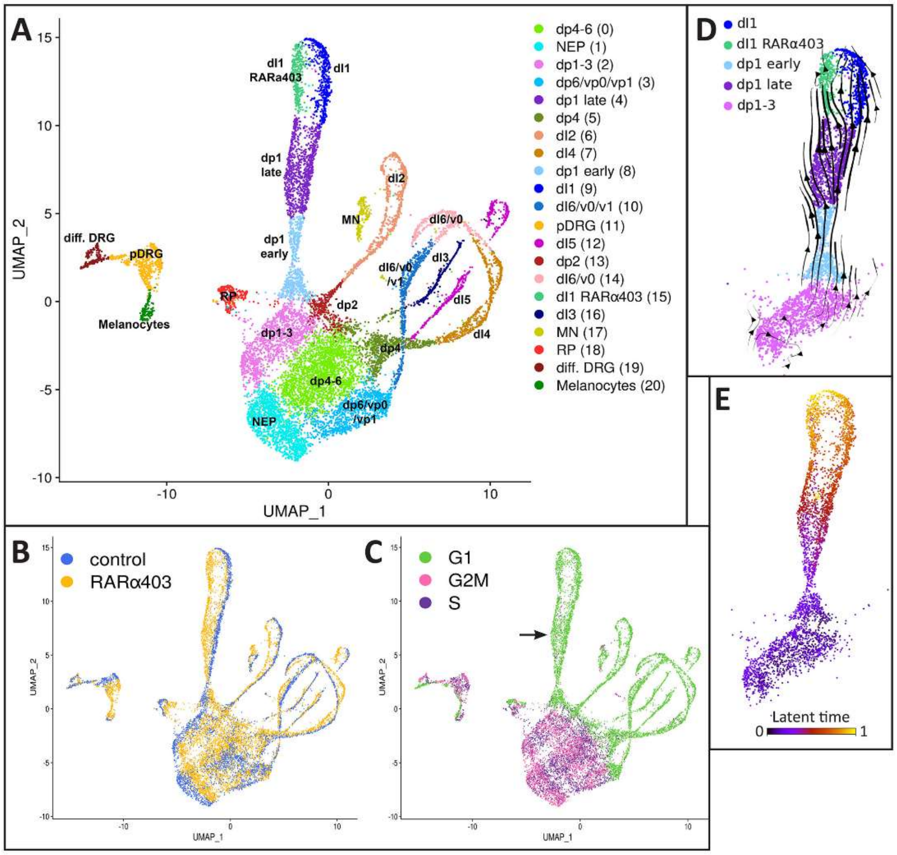


A) UMAP of control and RARα403-treated samples, colored by cluster. Cluster numbers indicated in parentheses.

B) UMAP of entire dataset colored by sample.

C) UMAP of the combined control and RARα403-treated sample, colored by cell cycle phase. Mitotic progenitors are located in the main body of the UMAP with only few cycling cells along the treated arm, primarily on its left side corresponding to the RARα403-treated cells (arrow).

D) RNA velocity vectors projected on a UMAP of the dI1 clusters of the control and RARα403-treated samples.

E) Latent time analysis of the dI1 clusters of the entire dataset. The earliest time point is depicted in black/purple and the latest in yellow.

**Supplementary Figure 14. Inhibition of RA signaling alters gene expression patterns along the dI1 interneuron developmental trajectory**

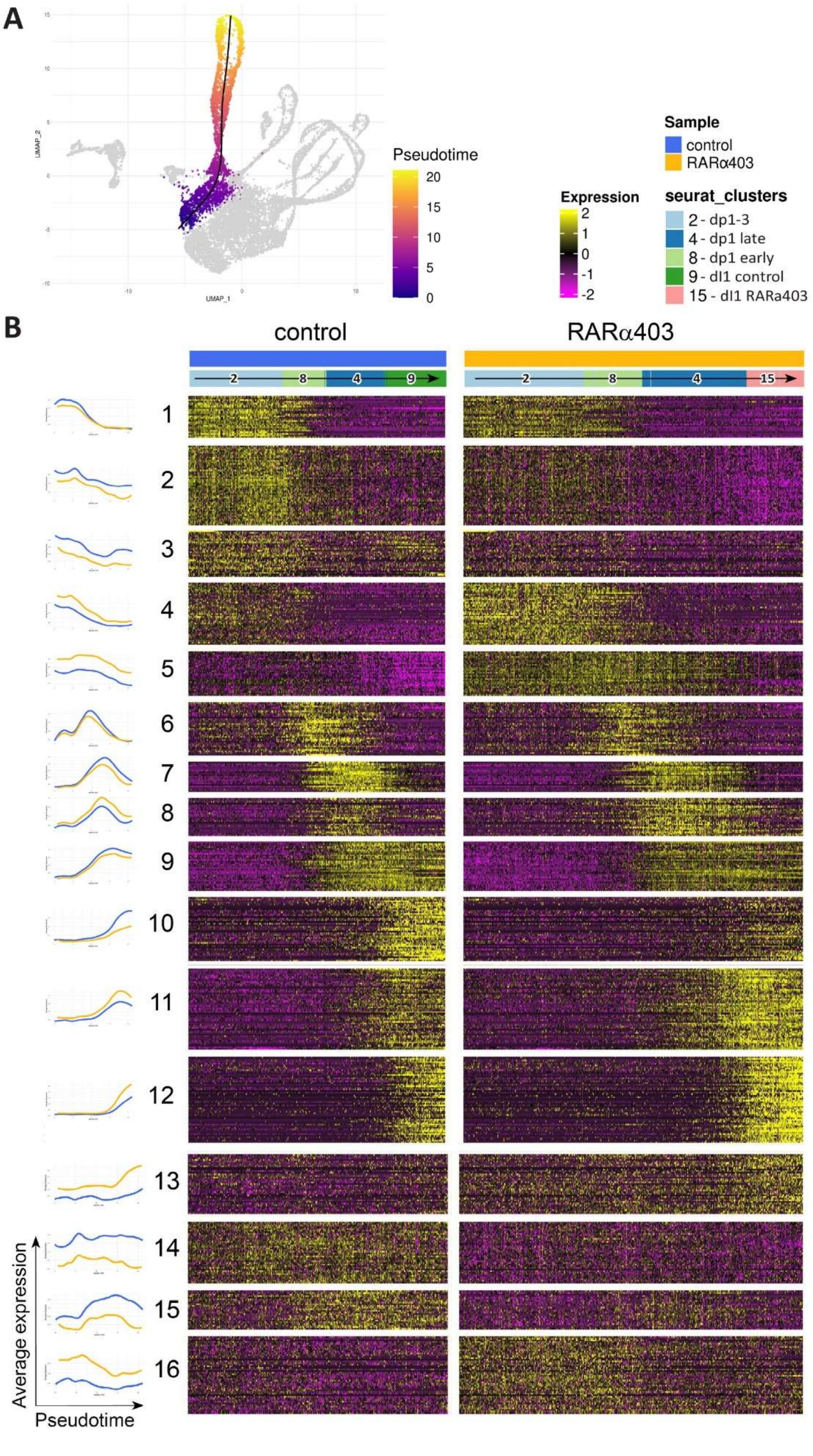

A) UMAP of the entire dataset. Trajectory inference with cluster 2 set as the starting cluster and cells of clusters 9 and 15 were set as the end clusters. The fitted principal curve is shown in black. The dI1 clusters are colored by pseudotime (Slingshot).

B) Comparative gene expression analysis between control and RARα403-treated samples. The left panel (line graphs) displays the average expression levels of each gene group along the pseudotime. A total of 571 genes showed significant differential expression (absolute fold change > 1.3 and adjusted p-value < 0.05) in at least one of the dI1 clusters. The scaled expression data were grouped into 16 groups using Euclidean distance. Group numbers shown on the left.

Groups 1-12- Differential expression was observed either between treated and control cells of clusters 2,8,4, or between cluster 9 (terminal dI1 control) and 15 (terminal dI1 treated).

Groups 13-16- Heatmap of differentially expressed genes along the trajectory in treated cells compared to control cells focusing on clusters with general expression patterns. Four groups that comprise 147 genes showed general expression pattern. The sample identity and cluster number are shown as top annotations.

**Supplementary Figure 15 – RA impacts the temporal regulation of Neuronal Differentiation and Signaling along the dI1 developmental trajectory**

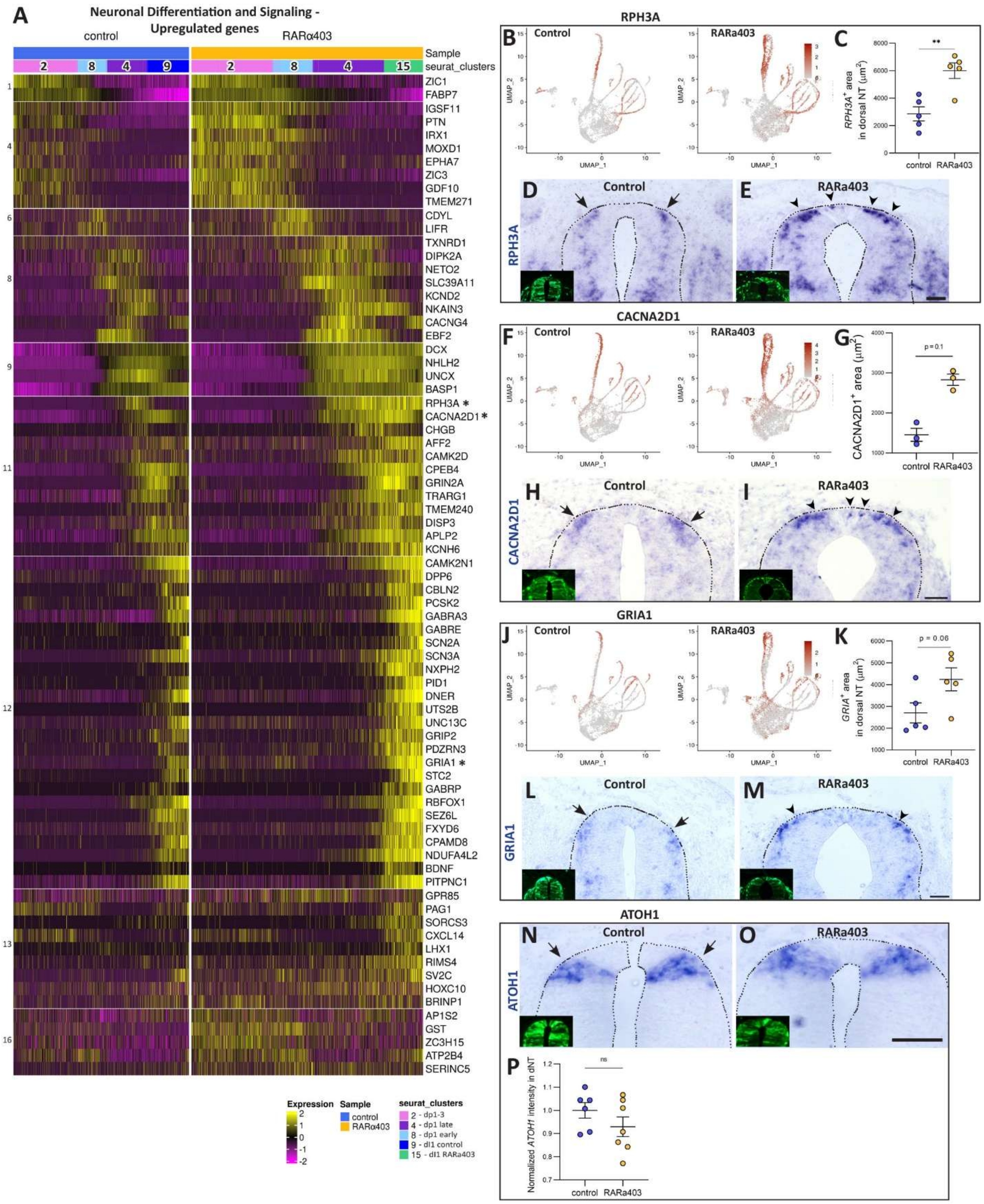


A) Heatmap of genes upregulated upon RA inhibition and associated with Neuronal Differentiation and Signaling. Cells are ordered by pseudotime and genes are classified according to the gene groups defined in Sup. Figure 14.

B-M) Validation of *RPH3A* (B-E), *CACNA2D1* (F-I) and *GRIA1* (J-M) expression changes (genes marked with asterisk in A). Embryos were electroporated with control PCAGG or RARα403 at E2.5, fixed at E4, and analyzed by ISH. Displayed are expression levels projected onto UMAPs (B,F,J), representative ISH images (D,E, H,I, L,M) and quantitative analyses (C,G,K).

B-E) *RPH3A* is expressed in intermediate stages under control conditions (B and arrows in D) but is upregulated and expands into the differentiating domain in treated embryos (B and arrowheads in E). Note presence of *RPH3A*$^{+}$ cells in the RP domain. **$p < 0.01$ by Student's t-test (N = 5 embryos per group, 15-20 sections per embryo).

F-I) *CACNA2D1* shows restricted expression in control embryos (H, arrows), while RARα403-treated embryos exhibit expanded expression in the differentiating dI1 domain (I, arrowheads). Note the presence of *CACNA2D1*$^{+}$ cells in the RP domain. Mann-Whitney test was applied (N = 3 embryos per group, 17-68 sections per embryo).

J-M) *GRIA1* shows restricted expression in control embryos (L, arrows), while RARα403-treated embryos exhibit enhanced and expanded expression in the differentiating dI1 domain (M, arrowheads). Student's t-test was applied (N = 5 embryos per group, 12-20 sections per embryo). Insets in D,E,H,I, and L,M illustrate the corresponding electroporated NTs.

N-P) ISH showing *ATOH1* expression in dI1 interneurons of a control (N, arrows) and a treated (O) sample. (P) No significant (ns) effect of RA deprivation on the intensity of *ATOH1* transcription. Student's t-test was applied (N = 6 and 7 embryos in control and treated samples, 12-20 sections per embryo). Insets in N, O illustrate the corresponding electroporated NTs. Scale bar, 50 μm.

220 **Supplementary Figure 16 – Retinoic acid signaling is required for correct positioning of**
221 **dI3 interneurons and regulation of dI3 and motoneuron numbers**

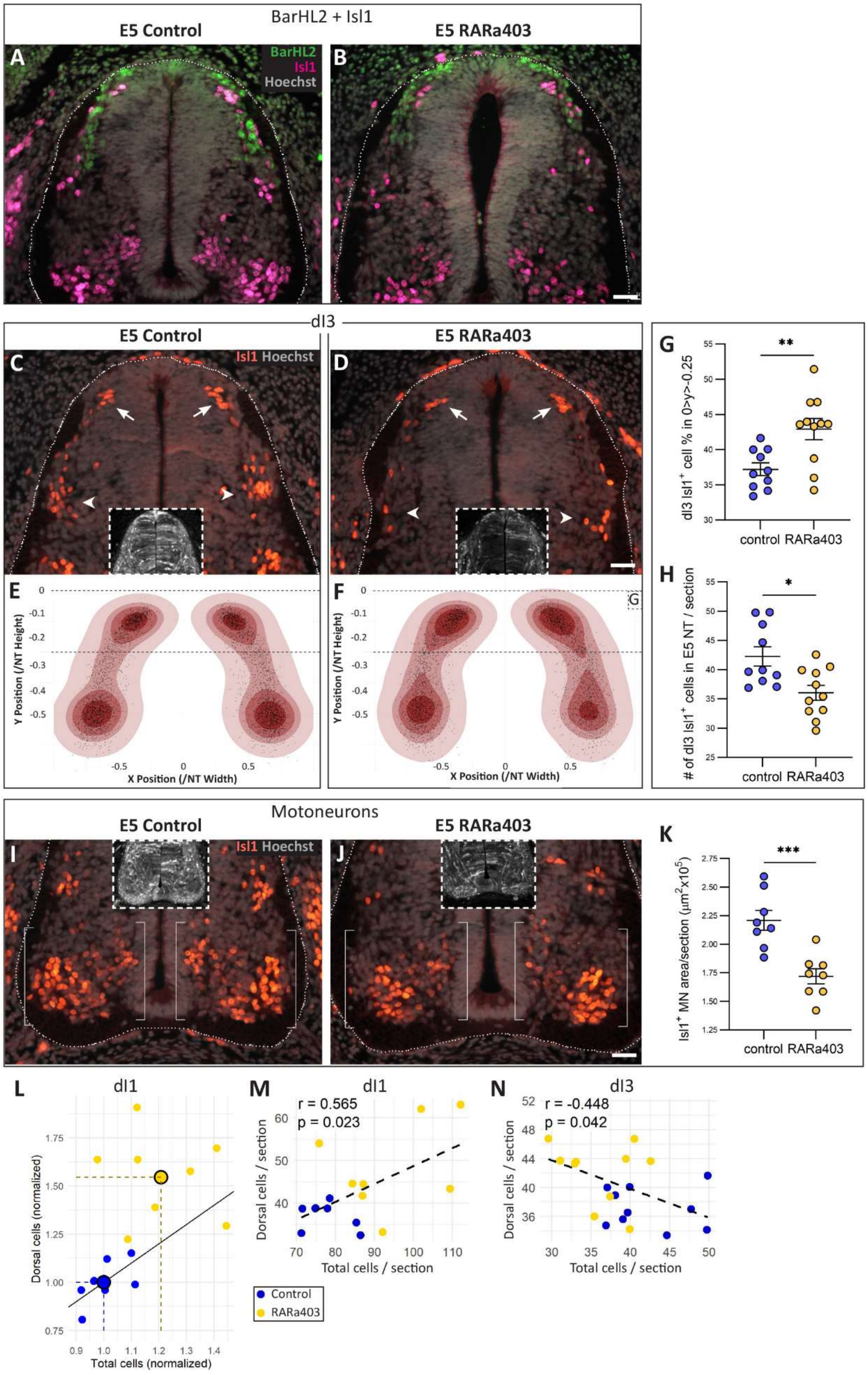


222

A-B) Embryos were electroporated at E2.5 with either control PCAGG (A) or RARα403 (B) plasmids, fixed at E5, and immunostained for BarHL2 (green) and Isl1 (magenta). Note that Isl1 marks both dorsal dI3 cells and ventral motoneurons.

C-D) Representative spinal cord sections showing the distribution of $Isl1^+$ dI3 interneurons. Indicated by arrows is the dorsal origin of dI3 interneurons, and by arrowheads the ventral destination domain.

E-F) Corresponding 2D density maps reveal altered distribution of dI3 interneurons in RARα403-treated embryos. Contours indicate increasing cumulative percentages of the total cell population. Black dots indicate individual cell positions. Dashed lines demarcate the dorsal region quantified in G. N = 5073 cells from 10 embryos (control) and 4759 cells from 11 embryos (RARα403); 15 sections per embryo. The same embryos and sections analyzed for dI1 distribution were used here, with a few additional embryos included to account for the lower abundance of dI3 interneurons compared to dI1.

G) Proportion of $Isl1^+$ cells retained within the dorsal 25% of the NT. Mann-Whitney test was applied; $^{**}p < 0.01$.

H) Total number of $Isl1^+$ dI3 cells at E5. A significant increase is observed in RARα403-treated embryos. Mann-Whitney test was applied; $^{*}p < 0.05$.

I-J) Representative spinal cord sections showing the distribution of $Isl1^+$ motoneurons, with brackets indicating their position relative to the dorsal $Isl1^+$ dI3 population.

K) Quantification of motoneuron area, showing a decrease in RARα403-treated embryos. Mann-Whitney test was applied; $^{***}p < 0.001$.

L) Comparison of normalized total $BarHL2^+$ and of dorsally located (within the region indicated in Fig. 8D-E) cell counts for control and RARα403-treated dI1 neurons. Each point represents an individual embryo. The diagonal line indicates the identity line (dorsal = total); embryos positioned above the line exhibit a higher proportion of dorsal cells relative to total. Large black-rimmed circles denote group means, showing 1.2-fold increase in total cell number and a 1.54-fold increase in dorsal cell number in RARα403-treated embryos compared to controls.

M-N) Correlation between total dI1 (M) or dI3 (N) cell number and the number of dorsally located cells in each population. The dorsal subsets were defined within the regions analyzed in Fig. 8D-E (dI1) and Sup. Fig. 18F-G (dI3). Each point represents an individual embryo. Dashed lines indicate the linear regression fit. A significant opposite correlation pattern was observed: total and dorsal cell numbers were positively correlated in dI1 neurons, but negatively correlated in dI3 neurons.

Scale bar, 50 μm.

**Supplementary Table S1- Quantitative data analysis**

Excel file with raw data and quantifications of relevant experiments
